# Formation of sulfoquinovosyl diacylglycerol by acylation of sulfoquinovosyl glycerol

**DOI:** 10.1101/2022.08.18.504050

**Authors:** Jessica Y. Cuevas-Rivas, Diego Rodriguez-Terrones, Napoleón González-Silva, Ángeles Moreno-Ocampo, Gabriela Guerrero, Ed Bergström, Jane E. Thomas-Oates, Otto Geiger, Isabel M. López-Lara

## Abstract

Sulfoquinovosyl diacylglycerol (SQDG) is a membrane-forming lipid present in photosynthetic organisms as well as in distinct bacteria growing in phosphate-limited environments. Four genes for SQDG biosynthesis were previously identified in *Rhodobacter sphaeroides*, the operon *sqdBDC* and *sqdA*. In this work, we found that SMc02490 of *Sinorhizobium meliloti* is an SqdA orthologue. Expression of the *S. meliloti sqdBDC* operon in *Escherichia coli* results in formation of sulfoquinovosyl glycerol (SQGro), while co-expression of this operon together with *smc02490* (*sqdA)* results in formation of SQDG. Furthermore, SqdA allows for incorporation of exogenous SQGro into SQDG in *S. meliloti* and in *E. coli* cultures, suggesting that presence of SqdA in bacteria permit them to use environmental SQGro for the biosynthesis of the membrane lipid SQDG. An *in vitro* enzymatic assay for the acyltransferase SqdA was developed. Cell-free crude extracts of *E. coli* expressing *sqdA* can efficiently convert [^35^S]-sulfoquinovosyl monoacylglycerol into SQDG using as acyl donor acyl carrier protein. Bioinformatic analyses reveal that this sulfoquinovose acylation pathway for SQDG biosynthesis is delimited to the *Hyphomicrobiales (Rhizobiales)* and *Rhodobacterales* orders of Alphaproteobacteria.

## Introduction

Sulfoquinovosyl diacylglycerols (SQDGs) are ubiquitous anionic glycolipids which are among the most abundant sulfur-containing bioorganic compounds. It is estimated that half of the sulfur in the biosphere is present within SQDG, on a scale similar to the amino acids cysteine and methionine (Goddard-Borger and Williams, 2017 and references therein). SQDGs are found in the membranes of all photosynthetic organisms, including higher plants, ferns, mosses, algae, dinoflagellates, diatoms and most photosynthetic bacteria (reviewed in Goddard-Borger and Williams, 2017). However, SQDG is also present in the membranes of some non-photosynthetic bacteria such as members of the *Rhizobiaceae* family (Cedergren and Hollingswoth, 1994), some Caulobacteria (Abraham *et al*., 1997) as well as in some Gram-positive bacteria (Sprott *et al*., 2006, Alcaraz *et al*., 2008). Because they are present in thylakoid membranes, SQDGs were thought to be essential for photosynthesis. However, requirement of SQDG for photosynthesis and growth varies across species (Kobayashi, 2016). Clearly, SQDG has a role as surrogate for phospholipids when their producers grow under phosphate limitation. Substitution of phosphatidylglycerol (PG) by SQDG has been described for different organisms such as *Synechococcus*, *Chlamydomonas* or *Arabidopsis* (Frentzen, 2004). Furthermore, some microorganisms living in oligotrophic environments such as the cyanobacterium *Prochlorococcus*, and the Gram-positive bacteria *Marinococcus*, *Salinicoccus*, and *Bacillus coahuilensis* contain SQDG as the major membrane lipid (Alcaraz *et al*., 2008; Sprott *et al*., 2006; Van Mooy *et al*., 2006). Although only demonstrated for the alga *Chlamydomonas reinhardtii*, SQDG has been proposed as a sulfur source for protein synthesis during sulfur limitation (Sugimoto *et al*., 2007).

The first genes for SQDG biosynthesis were discovered 30 years ago in *Rhodobacter sphaeroides*. Mutants in SQDG biosynthesis were identified by thin layer chromatography (TLC) from chemically mutagenized cells. After genetic complementation by introducing wild type cosmids, the *sqdA* gene was identified (Benning and Somerville, 1992a) as well as two genes that were part of a three gene operon in the order *sqdB*, ORF2, *sqdC* (Benning and Sommerville, 1992b). A fourth gene for SQDG biosynthesis (*sqdD*) was identified by insertional inactivation of the ORF2 flanked by the *sqdB* and *sqdC* genes (Rossak *et al*., 1995). The *sqdB* gene from *R. sphaeroides* was used to identify the first cyanobacterial *sqdB* gene in *Synechococcus* (Güler *et al*., 1996) while the cyanobacterial *sqdB* gene was used to identify the orthologous gene in *Arabidopsis* which was named SQD1 (Essigman *et al*., 1998). The *sqdB* gene of *Synechococcus* was found in an operon with another gene required for SQDG biosynthesis that was named *sqdX* (Güler *et al*., 2000). The protein encoded by *sqdX* has similarities to lipid glycosyltransferases but has no homology with the proteins encoded by *sqdA*, *sqdD* or *sqdC* genes identified in *R. sphaeroides*. Using the cyanobacterial *sqdX* gene was possible to isolate the plant orthologue SQD2 (Yu *et al*., 2000).

Genetic data show that the protein SqdB (SQD1 in plants) is the only one that has orthologues is all SQDG producing organisms (Benning *et al*., 2008). The recombinant SQD1 protein from *Arabidopsis* was shown to convert *in vitro* UDP-glucose and sulfite to UDP-sulfoquinovose (Sanda *et al*., 2001), thereby defining the initial step for SQDG biosynthesis (Fig. 1). In cyanobacteria and plants, SqdX/SQD2 transfer the sulfoquinovose moiety from UDP-sulfoquinovose to diacylglycerol (DAG) resulting in the formation of SQDG (Yu *et al*., 2002) (Fig. 1). Much less is known of the biochemical functions of the other SqdA, SqdD and SqdC proteins of *R. sphaeroides*. In agreement with the function of SqdB, the specific inactivation of *sqdD* led to the accumulation of UDP-sulfoquinovose (Rossak *et al*., 1995). SqdD shows homology with glycogenin, an autoglycosylating glycosyltransferase, and has no homology with the lipid glycosyltransferases SdqX or SQD2. Inactivation of *sqdC* led to the accumulation of sulfoquinovosyl-1-*O*-dihydroacetone (Rossak *et al*., 1997). This result suggests that SqdD transfers sulfoquinovose from UDP-sulfoquinovose to the possible acceptor dihydroxyacetone or dihydroxyacetone phosphate (Fig. 1). SqdC shows homology to small molecule reductases and could reduce the ketone group of the dihydroxyacetone residue to glycerol leading to the formation of sulfoquinovose glycerol (SQGro) as shown in Fig. 1. SqdA has homology to bacterial acyltransferases and could be involved in acylation of one or both hydroxyl groups of SQGro (Fig. 1). Such a pathway was proposed by Tamot and Benning (2009) but, so far, remains speculative. In any case, the previous results show that SQDG is assembled in *R. sphaeroides* by a route different from that known from plants and cyanobacteria (Fig. 1).

**Figure 1.**
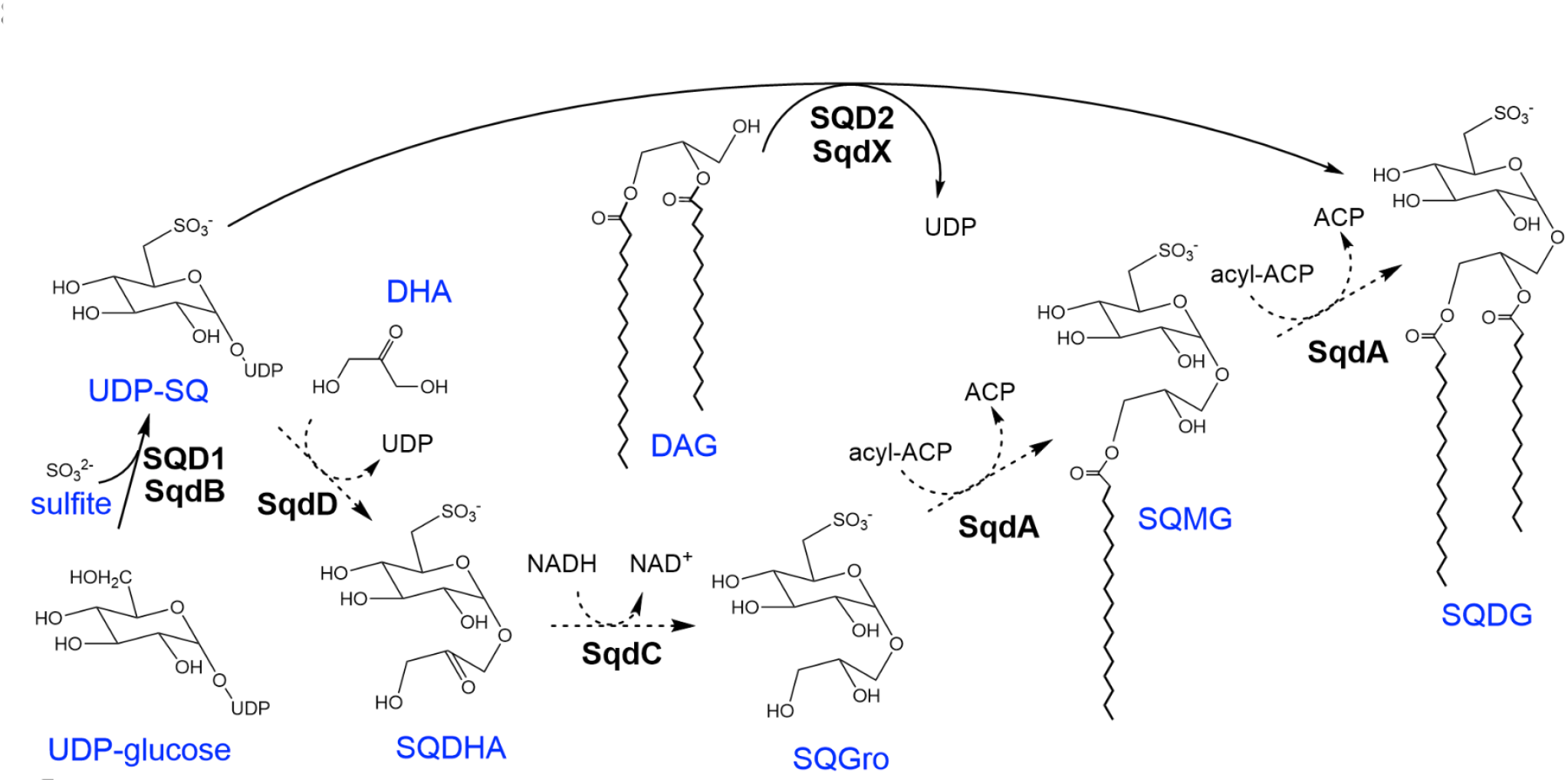
Pathways for SQDG biosynthesis. In all organisms producing SQDG, biosynthesis starts with the assembly of UDP-sulfoquinovose (UDP-SQ) from UDP-glucose and sulfite using the enzyme SqdB or its orthologue SQD1. In cyanobacteria and plants, SqdX or SQD2 transfer the sulfoquinovose moiety from UDP-SQ to diacylglycerol (DAG) leading to sulfoquinovose diacylglycerol (SQDG) formation. The proposed pathway for SQDG biosynthesis in *R. sphaeroides* after formation of UDP-SQ is indicated with dashed lines. SqdD transfer sulfoquinovose from UDP-SQ to dihydroxyacetone (DHA) leading to sulfoquinovosyl dihydroxyacetone (SQDHA) formation. SqdC reduces the ketone group in SQDHA to form sulfoquinovosyl glycerol (SQGro). SqdA has homology to acyltransferases and could performed one or two of the acylations required to form SQDG via sulfoquinovosyl monoacylglycerol (SQMG).

The α-proteobacerium *Sinorhizobium meliloti* was found to contain an *sqdBDC* operon and the encoded proteins had high similarity to the proteins encoded by the *sqdBDC* operon of *R. sphaeroides* with percentages of identity of 79, 62 and 47 % for SqdB, SqdD and SqdC, respectively (Weissenmayer *et al*., 2000). However, an orthologue for SqdA in *S. meliloti* was not known. In this work we have identified SMc02490 as the orthologue of SqdA which is required for SQDG biosynthesis. Expression of the sinorhizobial genes *sqdBDC* and *sqdA* in *Escherichia coli,* a bacterium not forming SQDG, led to the formation of significant amounts of SQDG while expression of only the three gene operon led to the formation of SQGro. Furthermore, we show that presence of *sqdA* only in *S. meliloti* or *E. coli* is sufficient to convert exogenous SQGro into SQDG. Finally, using *in vitro* assays, we show that the acyltransferase SqdA uses acyl carrier protein (AcpP) as fatty acyl donor to acylate the lyso-form of SQDG, sulfoquinovosyl monoacylglycerol (SQMG). We propose to name this second route for SQDG biosynthesis as SQGro acylation pathway and it is limited to members of the families *Rhizobacteriaceae* and *Rhodobacteraceae*.

## Results and discussion

### smc02490 is required for sulfoquinovosyl diacylglicerol biosynthesis in Sinorhizobium meliloti

The operon *sqdBDC* involved in SQDG formation was previously identified in *S. meliloti* (Weissenmayer *et al*., 2000). However, there is no ORF annotated as coding for SqdA in the genome of this organism (https://iant.toulouse.inra.fr/bacteria/annotation/cgi/rhime.cgi). Employing as query the SqdA sequence from *Rhodobacter sphaeroides* (Benning and Somerville, 1992a) against the *S. meliloti* proteome we identified SMc02490 that showed 39.3 % identity to SqdA in an overlap of 88 % of the amino acids of both proteins (E value of 1.25 x 10^-53^). SMc02490 is annotated as conserved hypothetical protein, but previous microarray data have shown that expression of *smc02490* is induced under phosphate limitation via the transcriptional factor PhoB (Krol and Becker, 2004). In agreement with this result, a *pho* box was identified 41 bp upstream from the annotated translational start codon of *smc02490* (Yuan *et al*., 2006). Analysis with the STRING software (http://string.embl.de) showed the highest level of coocurence of *smc02490* with the genes of the operon *sqdBDC*. Importantly, mutants of *S. meliloti* in *smc02490* showed a high cofitness value to each of the mutants in genes of the *sqdBDC* operon (values of 0.92, 0.89 and 0.85 for mutants in *sqdD*, *sqdB* and *sqdC*, respectively). This means that the mutants behaved in a very similar way in diverse growth experiments and suggests that they are probably involved in a same pathway (Price *et al*., 2018). All of these data support that SMc02490 is the sinorhizobial SqdA orthologue involved in SQDG biosynthesis and are in agreement with data indicating that SQDG formation in *S. meliloti* is increased in response to phosphate limitation controlled through the PhoB regulator (Geiger *et al*., 1999). A deletion mutant (NG8) was generated in which most of the ORF coding for SMc02490 had been replaced by a cassette conferring chloramphenicol resistance. Thin-layer chromatographic (TLC) analysis of radiolabeled polar lipids obtained from phosphate-limited cultures of different *S. meliloti* strains reveals that the slowest migrating spot present in the wild type strain 1021 is absent in the strain lacking *smc02490*. This same spot is missing in an *sqdB* mutant making it possible to be assigned to SQDG (Fig. 2). Expression of *smc02490* in *trans* in the mutant NG8 restores formation of SQDG while introduction of the empty vector does not (Fig. 2). The lipid profile of strain OD1 which is deficient in formation of DGTS and of OL, but still forms SQDG (López-Lara *et al*., 2005), was also analyzed in parallel in order to identify the migration of phosphorous-free membrane lipids (Fig. 2).

**Figure 2.**
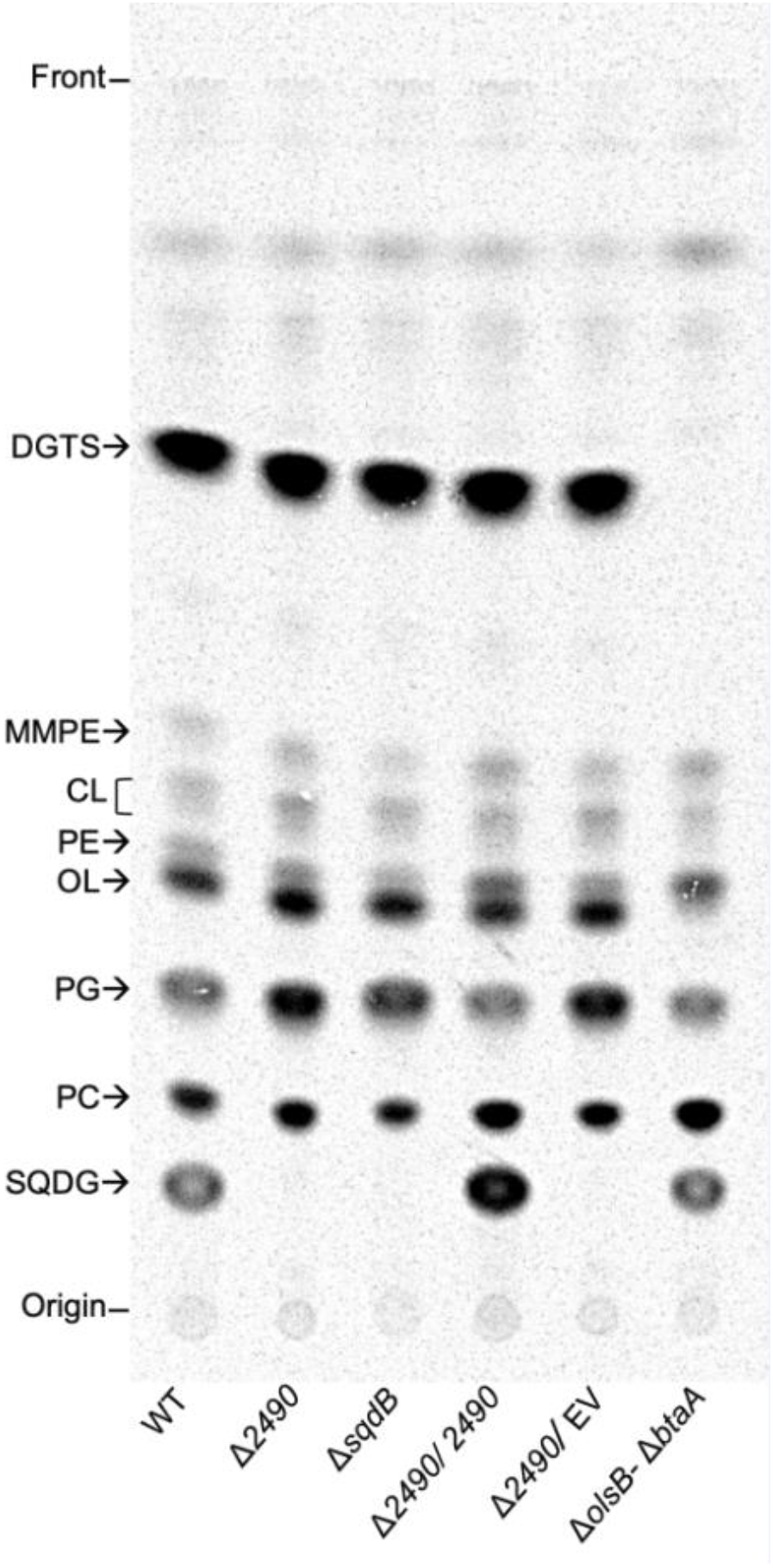
SMc02490 is essential for SQDG biosynthesis in *Sinorhizobium meliloti*. *S. meliloti* strains were radiolabeled during growth on Sherwood minimal medium with limiting (0.02 mM) concentrations of phosphate and lipids were separated by one-dimensional TLC. The strains analyzed were *S. meliloti* wild type 1021 (WT), and derivatives thereof: NG8 (Δ*smc02490*), SLD12 (Δ*sqdB*), NG8 carrying pDRT007 (Δ*smc02490*/*smc02490*), NG8 carrying the empty vector pBBR1MCS-2 (Δ*smc02490*/EV) and OD1 (Δ*olsB-ΔbtaA*). Migration of phosphorus-free lipids was identified by its absence in respective mutants: SQDG, sulfoquinovosyl diacylglycerol (absent in Δ*sqdB*); DGTS, diacylglyceryl trimethylhomoserine; and OL, ornithine lipid (both absent in Δ*olsB-ΔbtaA*). Migration of phosphorus-containing lipids PC, phosphatidylcholine; PG, phosphatidylglycerol; PE, phosphatidylethanolamine, CL, cardiolipin; MMPE, monomethylphosphatidylethanolamine were assigned by comparing different mutant strains (Fig. S1).

In a wild type strain grown under phosphate-limiting conditions, PG comprises 8 % and SQDG another 8 % of the total polar lipids while in an *smc02490* deficient mutant PG amounts to 16 % of the total lipids (Table 1). In different strains lacking SQDG, the amount of PG is increased while in the strain where SQDG is overproduced by expression of *smc02490* from plasmid pDRT007, the amount of PG is decreased. Except for the special case of mutant OD1, the total amount of PG plus SQDG is maintained in a similar range being between 13 and 20 % of the total polar lipids (Table 1). Therefore, quantification of polar lipids shows that in the absence of SQDG formation, the anionic lipid PG is compensating for it. Both PG and SQDG are anionic glycerolipids and this mutual compensation has also been described in different photosynthetic organisms such as *Synechococcus*, *Chlamydomonas* or *Arabidopsis*. Therefore, under different conditions, the combined PG and SQDG content remains largely constant (reviewed in Frentzen, 2004). To demonstrate the chemical identity of the SQDGs, lipids were isolated from the mutant deficient in *smc02490* carrying the empty vector pBBR1MCS-2 and from the mutant strain complemented *in trans* with the broad-host plasmid pBBR1MCS-2 carrying *smc02490* (plasmid pDRT007). Using matrix-assisted laser desorption/ionisation (MALDI) Fourier-transform ion cyclotron resonance (FT-ICR) mass spectrometry in negative ion mode, a total of nine *m/z* values consistent with SQDG species were observed in the spectrum of the lipid extract from the complemented mutant strain (Table 2), while these were not detected in the spectra of extracts obtained from the mutant carrying the empty vector. The accurate *m/*z signals provided using FT-ICR mass spectrometry are clearly consistent with elemental compositions of SQDGs (all *m/*z errors are well below 0.5 ppm). The two most intense signals were observed at nominal *m/z* values of 819 and 833; on high mass accuracy collision induced dissociation (CID) product ion analyses, ions were formed consistent with SQDG species carrying C16:0 with C18:1 for *m/z* 819 and C16:0 with C19:1 for *m/z* 833, on the basis of product ions formed by elimination of each of the fatty acids (Table 2). Taken together with the results of similar product ion analyses of the remaining seven signals the data showed that the full range of SQDG species identified (Table 2) is similar to those SQDGs identified in other studies using samples of *S. meliloti* grown under phosphate limitation (Geiger *et al*., 1999; Basconcillo *et al*., 2009a).

**Table 1.** Membrane lipid composition of *S. meliloti* strains under phosphate limitation.^a^

| Lípid | Composition (% of total <sup>14</sup>C) |  |  |  |  |  |
| --- | --- | --- | --- | --- | --- | --- |
| | WT | $\Delta 2490$ | $\Delta sqdB$ | $\Delta 2490/2490$ | $\Delta 2490/ EV$ | $\Delta olsB-\Delta btaA$ |
| DGTS | 57.3 ± 3.7 | 55.2 ± 8.3 | 59.9 ± 6 | 51.4 ± 6.8 | 51.1 ± 5.5 | ND |
| MMPE | 0.6 ± 0.4 | 0.6 ± 0.6 | 0.3 ± 0.5 | 0.8 ± 0.7 | 0.6 ± 0.6 | 2.4 ± 1.8 |
| CL | 3.1 ± 2.1 | 2.9 ± 1.7 | 3.2 ± 1.8 | 2.6 ± 1.4 | 3.5 ± 1.7 | 4 ± 2.7 |
| PE+OL | 14.8 ± 1.4 | 17.4 ± 2.1 | 17.3 ± 2.3 | 15.3 ± 1 | 20.5 ± 3.2 | 17.9 ± 3.9 |
| PG | 8.1 ± 2 | 16.2 ± 3.4 | 13.1 ± 3.4 | 5 ± 1 | 16.6 ± 3.9 | 11.7 ± 1.1 |
| PC | 8.2 ± 2 | 7.7 ± 3.1 | 6.2 ± 2.1 | 10.1 ± 3.4 | 7.7 ± 3.7 | 48.1 ± 1.6 |
| SQDG | 7.9 ± 3.2 | ND | ND | 14.8 ± 5.9 | ND | 15.9 ± 5.7 |
| SQDG+PG | 16 ± 4.1 | 16.2 ± 3.4 | 13.1 ± 3.4 | 19.8 ± 6.6 | 16.6 ± 3.9 | 27.6 ± 6.2 |

<sup>a</sup> Membrane lipid composition of *Sinorhizobium meliloti* 1021 wild type (WT), and derivatives of thereof: NG8 ( $\Delta 2490$ ), SLD12 ( $\Delta sqdB$ ), NG8 carrying pDRT007 ( $\Delta 2490/2490$ ), NG8 carrying the empty vector pBBR1MCS-2 ( $\Delta 2490/EV$ ) and OD1 ( $\Delta olsB-\Delta btaA$ ) after growth on low (0.02 mM) phosphate-containing minimal medium. The values shown are mean values ± standard deviation derived from three independent experiments. ND, not detected.

**Table 2.**
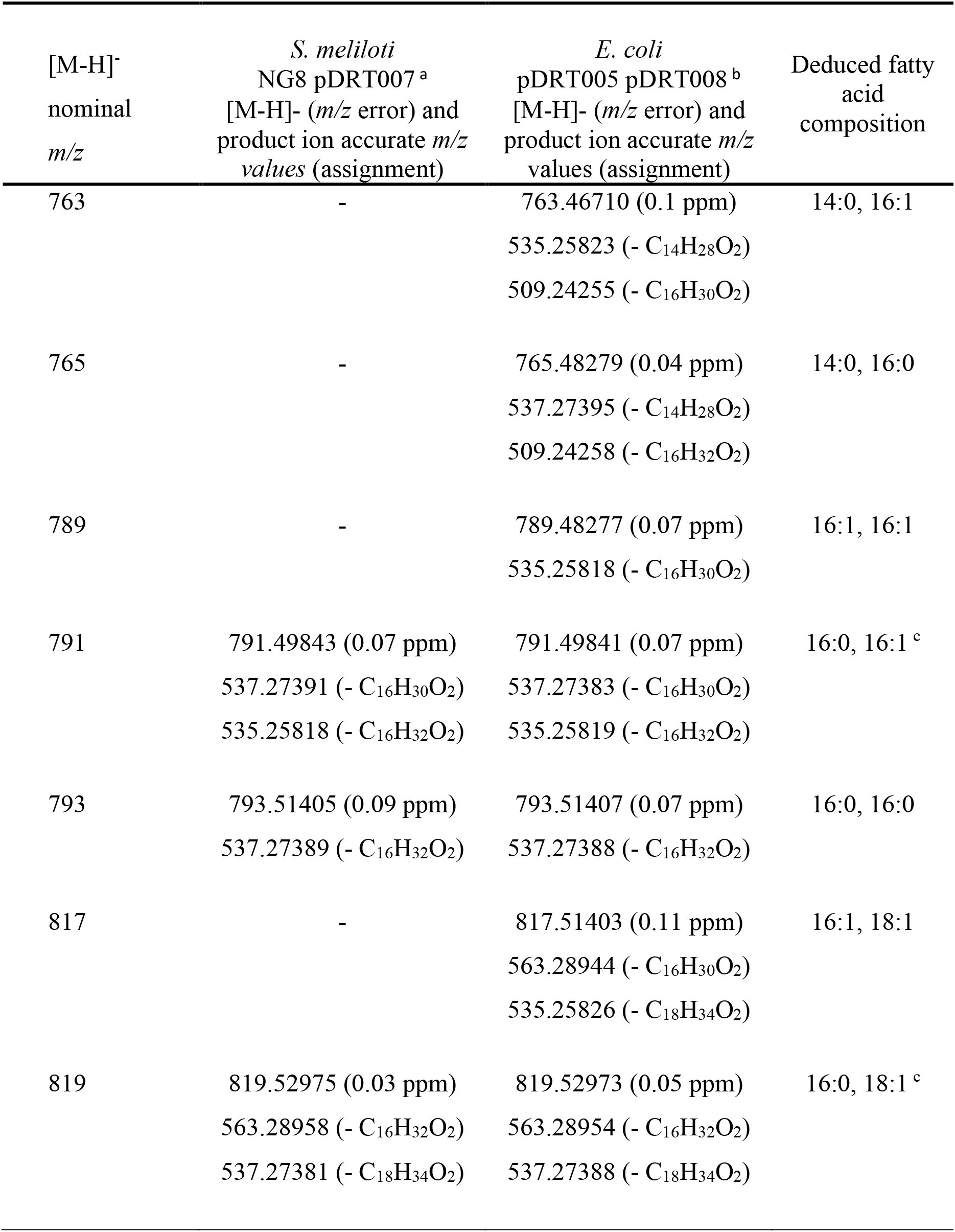
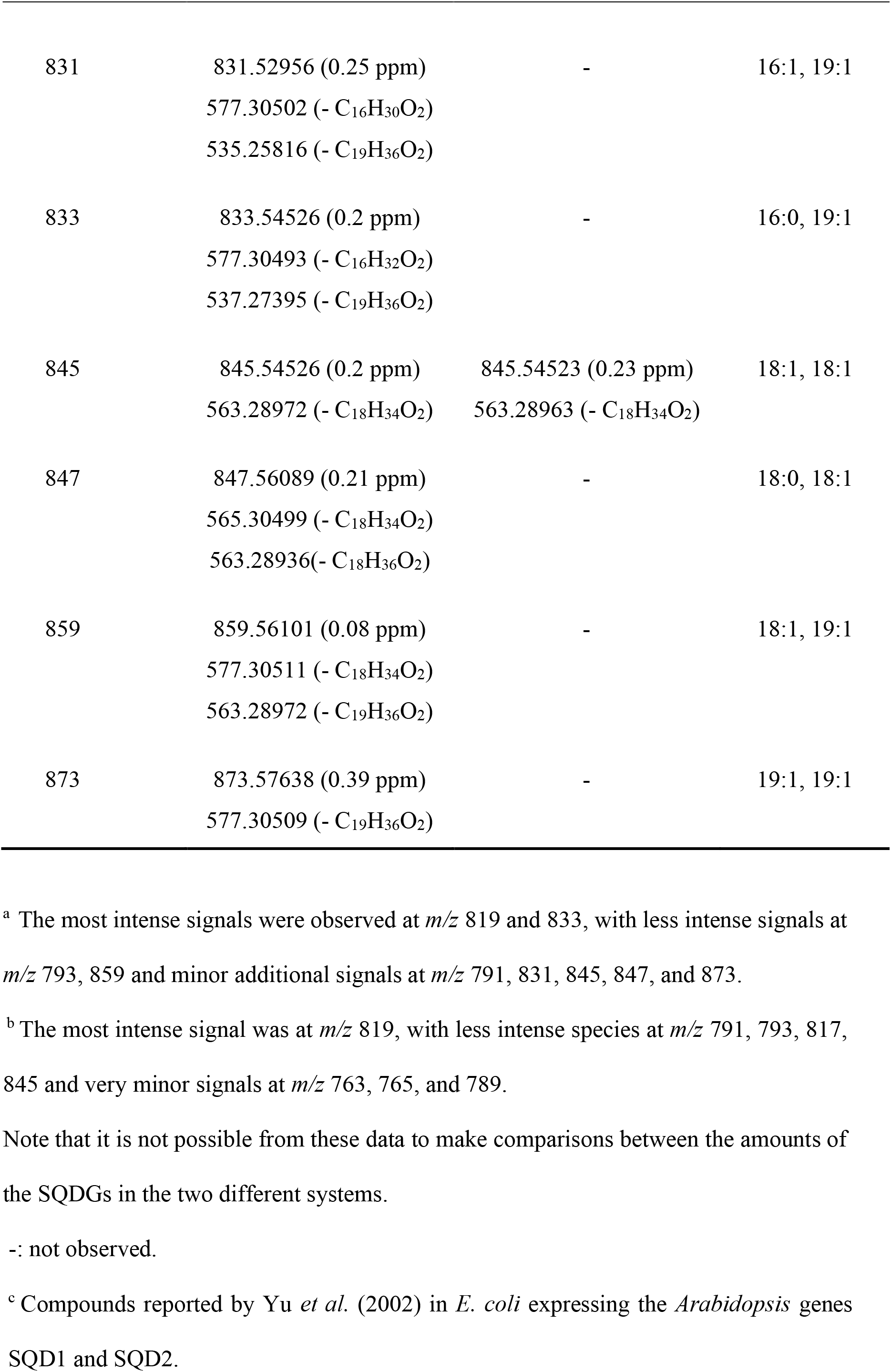
Mass spectrometric data from SQDG species in extracts of *S. meliloti* NG8 pDRT007 and *E. coli* BL21 (DE3) pLysS pDRT005 pDRT008.

| [M-H]<sup>-</sup><br>nominal<br>*m/z* | *S. meliloti*<br>NG8 pDRT007<sup>a</sup><br>[M-H]<sup>-</sup> (*m/z* error) and<br>product ion accurate *m/z*<br>values (assignment) | *E. coli*<br>pDRT005 pDRT008<sup>b</sup><br>[M-H]<sup>-</sup> (*m/z* error) and<br>product ion accurate *m/z*<br>values (assignment) | Deduced fatty<br>acid<br>composition |
| --- | --- | --- | --- |
| 763 | - | 763.46710 (0.1 ppm)<br>535.25823 (- C<sub>14</sub>H<sub>28</sub>O<sub>2</sub>)<br>509.24255 (- C<sub>16</sub>H<sub>30</sub>O<sub>2</sub>) | 14:0, 16:1 |
| 765 | - | 765.48279 (0.04 ppm)<br>537.27395 (- C<sub>14</sub>H<sub>28</sub>O<sub>2</sub>)<br>509.24258 (- C<sub>16</sub>H<sub>32</sub>O<sub>2</sub>) | 14:0, 16:0 |
| 789 | - | 789.48277 (0.07 ppm)<br>535.25818 (- C<sub>16</sub>H<sub>30</sub>O<sub>2</sub>) | 16:1, 16:1 |
| 791 | 791.49843 (0.07 ppm)<br>537.27391 (- C<sub>16</sub>H<sub>30</sub>O<sub>2</sub>)<br>535.25818 (- C<sub>16</sub>H<sub>32</sub>O<sub>2</sub>) | 791.49841 (0.07 ppm)<br>537.27383 (- C<sub>16</sub>H<sub>30</sub>O<sub>2</sub>)<br>535.25819 (- C<sub>16</sub>H<sub>32</sub>O<sub>2</sub>) | 16:0, 16:1<sup>c</sup> |
| 793 | 793.51405 (0.09 ppm)<br>537.27389 (- C<sub>16</sub>H<sub>32</sub>O<sub>2</sub>) | 793.51407 (0.07 ppm)<br>537.27388 (- C<sub>16</sub>H<sub>32</sub>O<sub>2</sub>) | 16:0, 16:0 |
| 817 | - | 817.51403 (0.11 ppm)<br>563.28944 (- C<sub>16</sub>H<sub>30</sub>O<sub>2</sub>)<br>535.25826 (- C<sub>18</sub>H<sub>34</sub>O<sub>2</sub>) | 16:1, 18:1 |
| 819 | 819.52975 (0.03 ppm)<br>563.28958 (- C<sub>16</sub>H<sub>32</sub>O<sub>2</sub>)<br>537.27381 (- C<sub>18</sub>H<sub>34</sub>O<sub>2</sub>) | 819.52973 (0.05 ppm)<br>563.28954 (- C<sub>16</sub>H<sub>32</sub>O<sub>2</sub>)<br>537.27388 (- C<sub>18</sub>H<sub>34</sub>O<sub>2</sub>) | 16:0, 18:1<sup>c</sup> |
| 831 | 831.52956 (0.25 ppm)<br>577.30502 (- C<sub>16</sub>H<sub>30</sub>O<sub>2</sub>)<br>535.25816 (- C<sub>19</sub>H<sub>36</sub>O<sub>2</sub>) | - | 16:1, 19:1 |
| 833 | 833.54526 (0.2 ppm)<br>577.30493 (- C<sub>16</sub>H<sub>32</sub>O<sub>2</sub>)<br>537.27395 (- C<sub>19</sub>H<sub>36</sub>O<sub>2</sub>) | - | 16:0, 19:1 |
| 845 | 845.54526 (0.2 ppm)<br>563.28972 (- C<sub>18</sub>H<sub>34</sub>O<sub>2</sub>) | 845.54523 (0.23 ppm)<br>563.28963 (- C<sub>18</sub>H<sub>34</sub>O<sub>2</sub>) | 18:1, 18:1 |
| 847 | 847.56089 (0.21 ppm)<br>565.30499 (- C<sub>18</sub>H<sub>34</sub>O<sub>2</sub>)<br>563.28936(- C<sub>18</sub>H<sub>36</sub>O<sub>2</sub>) | - | 18:0, 18:1 |
| 859 | 859.56101 (0.08 ppm)<br>577.30511 (- C<sub>18</sub>H<sub>34</sub>O<sub>2</sub>)<br>563.28972 (- C<sub>19</sub>H<sub>36</sub>O<sub>2</sub>) | - | 18:1, 19:1 |
| 873 | 873.57638 (0.39 ppm)<br>577.30509 (- C<sub>19</sub>H<sub>36</sub>O<sub>2</sub>) | - | 19:1, 19:1 |

<sup>a</sup> The most intense signals were observed at *m/z* 819 and 833, with less intense signals at *m/z* 793, 859 and minor additional signals at *m/z* 791, 831, 845, 847, and 873.
<sup>b</sup> The most intense signal was at *m/z* 819, with less intense species at *m/z* 791, 793, 817, 845 and very minor signals at *m/z* 763, 765, and 789.
Note that it is not possible from these data to make comparisons between the amounts of the SQDGs in the two different systems.
-: not observed.
° Compounds reported by Yu *et al.* (2002) in *E. coli* expressing the *Arabidopsis* genes SQD1 and SQD2.

All data presented in this section demonstrate that SMc02490 is required for SQDG biosynthesis in *S. meliloti* and that it is homologous to the SqdA ORF originally identified in *Rhodobacter sphaeroides* (Benning and Somerville, 1992a). Therefore, we propose SMc02490 be renamed as SqdA.

### Expression of sqdA and sqdBDC from plasmids in S. meliloti significantly increases SQDG formation under conditions of sufficient phosphate

The *S. meliloti* mutant NG8 carrying in *trans* the gene *sqdA* significantly increases the amount of SQDG formed with respect to the amount formed by the wild type strain under conditions of phosphate limitation (14.8 ± 5.9 vs 7.9 ± 3.2, Table 1), indicating a dose effect due to the augmented level of *sqdA* provided by the plasmid. To study the effect provided by extra copies of the *sqdBDC* operon in an *S. meliloti* background, we constructed a pRK404-derived plasmid carrying the *sqdBDC* genes (see material and methods). Under conditions of phosphate limitation, the presence of pRK404 carrying *sqdBDC* did not significantly increase the formation of SQDG (Fig. 3A, Table S1). However, at the same growth conditions, the strain carrying pBBR1MCS-2 with *sqdA* doubled the amount of SQDG formed with respect to the strain carrying two empty vectors (Fig. 3A, Table S1). This could be due to the fact that the pBBR1-derived plasmids are of higher copy number than the low copy number IncP-plasmid pRK404 (Kovach *et al*., 1995). Coexpression of *sqdA* and *sqdBDC* from both broad host range plasmids (pDRT007 and pDRT008) in the wild type *S. meliloti* under phosphate limiting conditions did not lead to a significant increase of SQDG formation with respect to a wild type strain carrying only extra copies of *sqdA* (Fig. 3A, supplementary Table S1, Fig. S2).

**Figure 3.**
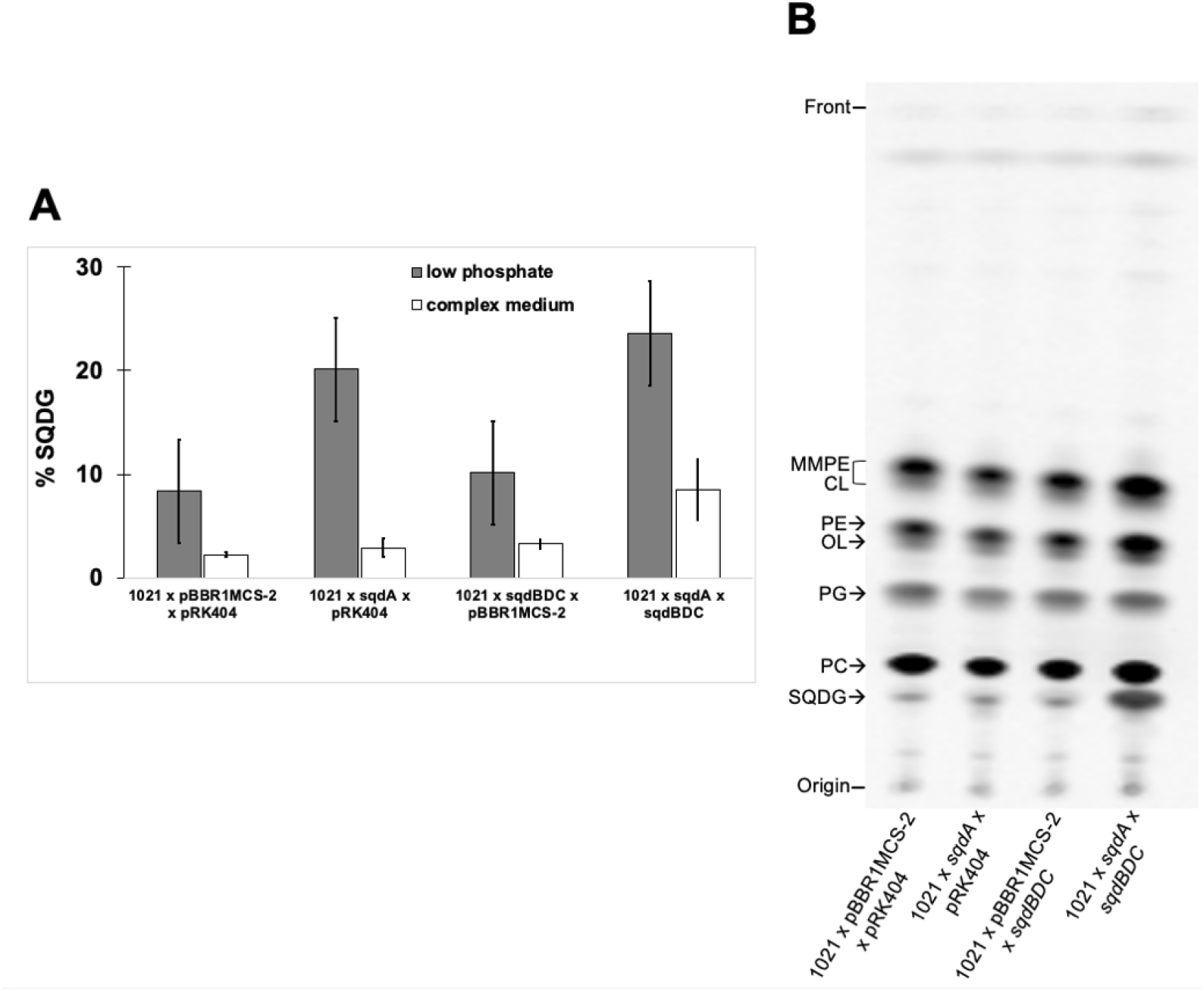
Expression of *sqdA* and *sqdBDC* from plasmids significantly increases the amount of SQDG formed in *S. meliloti*. **A.** Percentage of SQDG formed by *S. melioti* 1021 wild type carrying either two empty vectors (1021 x pBBR1MCS-2 x pRK404), pDRT007 and pRK404 (1021 x *sqdA* x pRK404), pDRT008 and pBBR1MCS-2 (1021 x *sqdBDC* x pBBR1MCS-2) or pDRT007 and pDRT008 (1021 x *sqdA* x *sqdBDC*) grown on Sherwood minimal medium with limiting (0.02 mM) concentrations of phosphate (grey bars) or on PYCa^++^ complex medium (white bars). Bars and error bars represent the mean and standard deviation of the SQDG percentage obtained for each strain from there independent experiments. A representative TLC of the lipid profile from the different strains grown on low phosphate medium is shown in Supporting information Fig. S2 and of the lipid profile of the strains grown on complex medium is shown in (B). **B**. Separation on TLC of ^14^C-labeled lipids formed by *S. melioti* 1021 wild type carrying either two empty vectors (1021 x pBBR1MCS-2 x pRK404), pDRT007 and pRK404 (1021 x *sqdA* x pRK404), pDRT008 or pBBR1MCS-2 (1021 x *sqdBDC* x pBBR1MCS-2) or pDRT007 and pDRT008 (1021 x *sqdA* x *sqdBDC*) grown on PYCa^++^ complex medium.

TLC analyses showed that the wild type strain *S. meliloti* carrying *in trans* extra copies of *sqdA* or of *sqdBDC* grown on PYCa^++^ complex medium produced similar amounts of SQDG as the wild type carrying two empty vectors. However, the wild type strain *S. meliloti* 1021 carrying plasmids with *sqdA* and with *sqdBDC* grown on PYCa^++^ complex medium produced a significantly higher amount of SQDG (8.5 ± 2.9 % of the total lipids) than strains containing only extras copies of *sqdA* or *sqdBDC* (Fig. 3B, Table S1). Again, these results showed that *sqdA* works together with *sqdBDC* for SQDG formation.

### Expression of sinorhizobial sqdA and sqdBDC in Escherichia coli results in SQDG formation

Biosynthesis of SQDG in *Arabidopsis* proceeds in two steps (Fig. 1) and expression of SQD1 and SQD2 in *E. coli* led to SQDG formation in this bacterium (Yu *et al*., 2002). Since homologous expression of *sqdA* and the operon *sqdBDC* led to SQDG formation in *S. meliloti* when grown in complex medium, we explored whether heterologous expression of the four genes in *E. coli* would lead to SQDG formation. As expected, expression of only *sqdA* in *E. coli* did not lead to formation of a compound with a mobility similar to SQDG. However, expression of *sqdA* and *sqdBDC* in *E. coli* led to a prominent spot that shows similar mobility to SQDG formed in *S. meliloti* (Fig. 4A). We used two compatible vectors, a pET vector and pRK404, and we studied the lipid profile obtained from different combinations of plasmids in the *E. coli* BL21(DE3) pLysS strain. No new bands with similar mobility to SQDG were visible in lipid extracts from *E. coli* cells containing two empty vectors, or strains that were carrying *sqdA* or the *sqdBDC* operon in combination with empty vectors. However, when *E. coli* cells contained constructs carrying *sqdA* (pDRT005) and *sqdBDC* (pDRT008) a new compound appeared, which shows similar mobility to SQDG in the lipid extract of *S. meliloti* (Fig. 4A and B).

**Figure 4.**
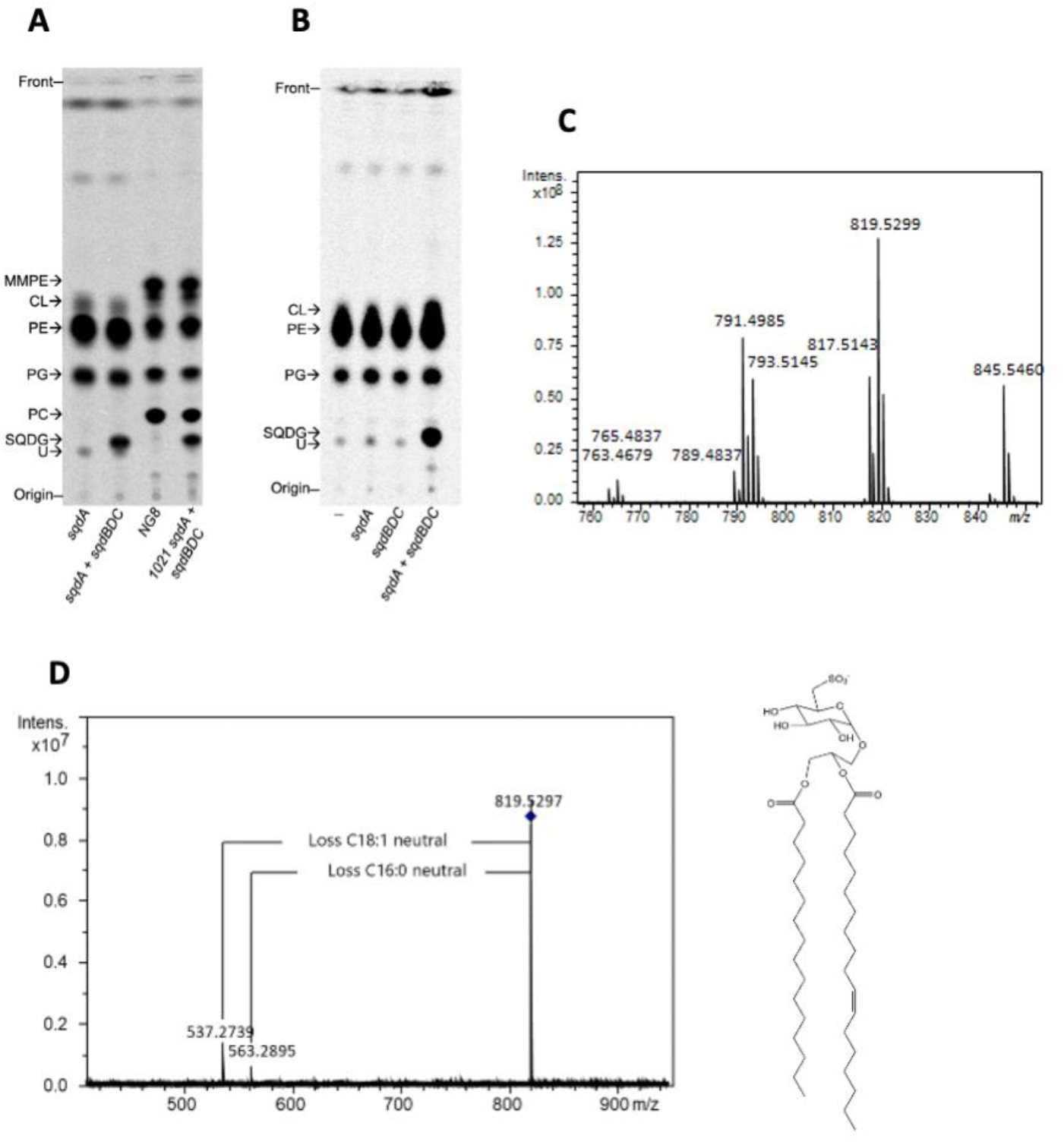
Expression of sinorhizobial *sqdBDC* and *smc02490* (*sqdA)* in *Escherichia coli* results in SQDG formation. **A.** ^14^C-acetate labeled lipids from *E. coli* BL21(DE3) pLysS containing either pDRT005 (*sqdA*) or pDRT005 plus pDRT008 (*sqdA + sqdBDC*) and lipids form *S. meliloti* deficient in *sqdA* (NG8) or from *S. meliloti* wild type carrying pDRT007 and pDRT008 (1021 *sqdA* + *sqdBDC*). **B.** ^14^C-acetate labeled lipids from *E. coli* BL21(DE3) pLysS containing either the empty vectors pET17b and pRK404 (-), pRK404 and pDRT005 (*sqdA*), pET17b and pDRT008 (*sqdBDC*) or pDRT005 plus pDRT008 (*sqdA + sqdBDC*). **C.** Partial MALDI mass spectrum of lipids from *E. coli* BL21(DE3) pLysS containing pDRT005 plus pDRT008 showing different SQDG signals. **D.** Collision induced dissociation product ion spectrum of the ion at *m/z* 819 produces only two very clear product ions derived by loss of each of the two fatty acyl chains. Tentative structure shown at the right of the panel. Abbreviations for lipids spots in TLCs as in Fig. 2. U: unknown lipid.

Lipids for mass spectrometric analysis were isolated from each of the four *E. coli* strains containing either the empty vectors pET17b and pRK404, pRK404 and pDRT005 (*sqdA*), pET17b and pDRT008 (*sqdBDC*) or pDRT005 and pDRT008 (*sqdA + sqdBDC*). Using MALDI FT-ICR mass spectrometry in negative ion mode, a total of eight *m/z* values consistent with SQDG species were observed only in the spectrum of the lipid extract from the *E. coli* strain carrying plasmids with *sqdA* and *sqdBDC* (Fig. 4C, Table 2), while these were not detected in the spectra of extracts obtained from any of the other three *E. coli* strains carrying either two empty vectors or combinations of vectors with only *sqdA* or the *sqdBDC* operon. The most intense signal, with a nominal *m/z* of 819 observed in the spectra of *E. coli* extracts was also one of the most intense signals observed in the spectra of extracts of *S. meliloti* (Table 2). The accurate *m/*z signals provided using FT-ICR mass spectrometry are clearly consistent with elemental compositions of SQDGs (*m/z* errors are well under 0.5 ppm). High mass accuracy CID product ion analysis of this species was consistent with SQDG carrying C16:0 and C18:1, on the basis of product ions formed by elimination of each of the fatty acids (Fig. 4D). Similar analyses of the remaining SQDG signals showed that the range of SQDG species identified in lipid extracts of *E. coli* carrying *sqdA* and *sqdBDC* is different from the range of SQDG species observed in extract of *S. meliloti* (Table 2). We detected SQDG species with C14:0 fatty acids in *E. coli* extracts which were not detected in those of *S. meliloti*. Notably, in *E. coli* extracts, SQDG species with the C19:1 fatty acid were not detected, while they are encountered in abundance in SQDG species from *S. meliloti*. Probably, the C19:1 fatty acid corresponds to lactobacillic acid, which is formed by cyclopropanation of C18:1. (Basconcillo *et al*., 2009b). Interestingly, expression of the plant genes SQD1 and SQD2 in *E. coli* also led to production of SQDG species with *m/z* values of 791 and 819 (Yu *et al*., 2002) which are the most abundant species detected in the strain carrying the sinorhizobial genes (Fig. 4C and Table 2). These results indicate that the SQDG species formed depend more on the host than on the pathway used.

In the *E. coli* strain carrying *sqdA* and *sqdBDC* grown on LB medium for 18 h after IPTG induction, the SQDG formed amounts to 15.5 ± 1.5 % of the total lipids, measured after ^14^C-acetate labeling. Therefore, we used this construct to obtain SQDG for further studies. This SQDG-producing *E. coli* strain could also be used to obtain SQDG or derivatives thereof for future biotechnological applications (Benning *et al*., 2008).

### Sulfoquinovosyl glycerol (SQGro) is an intermediary in the sinorhizobial pathway for SQDG biosynthesis

A significant amount of SQDG is formed after expression of *sqdA* and *sqdBDC* in *E. coli* (Fig. 4), indicating that the sinorhizobial enzymes for SQDG biosynthesis are functional in *E. coli*. Sulfoquinovose glycerol (SQGro) is proposed to be formed by action of the enzymes SqdB, SqdD and SqdC encoded by the *sqdBDC* operon (Fig. 1). SQGro can be easily obtained from SQDG since carboxylic acid ester linkages of glycerolipids are especially sensitive to treatment with mild alkali. As specified in materials and methods, non-labeled SQDG obtained from the overproducing *E. coli* strain, was treated with mild alkali and the putative SQGro was purified. Analysis of this sample by both MALDI and electrospray (ESI) FT-ICR mass spectrometry in negative ion mode resulted in detection of a signal consistent with SQGro. In the MALDI spectrum, this signal was of low intensity, at *m/z* 317.05490, an *m/z* accuracy for the elemental composition of SQGro of 0.3 ppm, while in the ESI spectrum, the equivalent signal was the base peak, at *m/z* 317.05481, an *m/z* accuracy of 0.07 ppm. Furthermore, the accurate *m/*z ESI product ion spectrum showed signals for the loss of the elements of glycerol (product ion at *m/z* 225.00742), the elements of glycerol and of water (ion at *m/z* 206.99686) and a further ion arising by cross ring cleavage of the sulfoquinovose monosaccharide, a typical carbohydrate fragmentation, arising by loss of the ring oxygen, and carbons 1 and 2 and their substituents (loss of C_5_H_12_O_5_, product ion at *m/z* 164.98630). After demonstrating that mild alkali hydrolysis of SQDG indeed led to SQGro formation, a control of this compound was prepared following the same procedure but from a culture grown in the presence of [^35^S]-sulfate. We made different plasmid constructs to be able to express genes in the order of the proposed biosynthetic pathway. ^35^S-labeled water-soluble compounds of the *E. coli* strain expressing *sqdB* contains a prominent spot not present in extracts of the *E. coli* strain carrying an empty vector (Fig. 5, Fig. S3). Based on its chromatographic behaviour and the known function of SqdB, we identified this compound as UDP-sulfoquinovose. Expression of *sdqBD* led to the accumulation of a faster migrating spot (Fig. 5). Rossak *et al*. (1997) found that an *R. sphaeroides* mutated in *sqdC*, but with intact *sqdBD*, accumulated sulfoquinovosyl-1-*O*-dihydroxyacetone. Furthermore, the faster migrating compound from the *E. coli* strain expressing *sqdBD* showed a similar R*_f_* as shown previously for sulfoquinovosyl dihydroxyacetone in an analogous TLC system analysis (Fig. 3 in Rossak *et al*., 1997). Expression in *E. coli* of plasmids carrying the genes *sqdBDC* led to the formation of a spot with identical migration to the prepared control SQGro (Fig. 5, Fig. S3). Also, the strain producing SQDG presents a spot with the same migration as SQGro (Fig. 5, Fig S3). We show here for the first time that SQGro is likely the product formed by the *sqdBDC* operon and reinforce the idea of SQGro as an intermediary in SQDG biosynthesis (Fig. 1).

**Figure 5.**
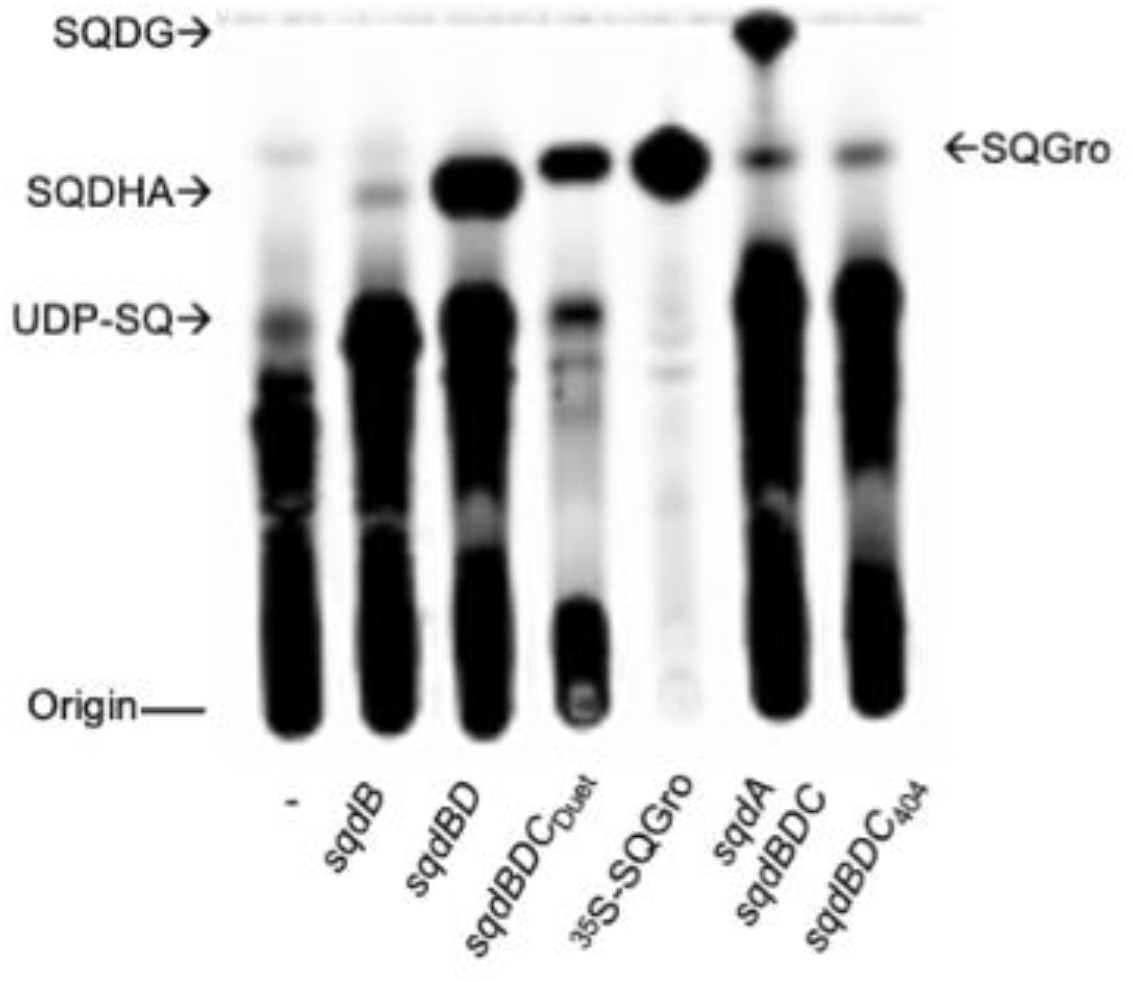
SQGro is an intermediary in the sinorhizobial pathway for SQDG biosynthesis. Thin layer chromatography of ^35^S-sulfate labelled, water-soluble compounds obtained from *E. coli* BL21(DE3) pLysS carrying either the empty vector pET17b (-), pJCR003 (*sqdB*), pJCR005(*sqdBD*), pJCR006 (*sqdBDC*_Duet_), pDRT005 and pDRT008 (*sqdA sqdBDC*), or pDRT008 (*sqdBDC*_404_) in comparison to ^35^S-labeled sulfoquinovosyl glycerol SQDG (^35^S-SQGro). prepared from purified ^35^S-SQDG. UDP-SQ: UDP sulfoquinovose, SQDHA: possible sulfoquinovosyl dihydroxyacetone. The mobile phase was ethanol: water: acetic acid (20:10:1, v/v).

### SqdA allows SQDG formation from exogenous SQGro

To further confirm SQGro as an intermediary of SQDG biosynthesis, we added ^35^S-SQGro to different *S. meliloti* strains. Addition of SQGro to *S. meliloti* wild type results in the detection of a labeled compound that migrated like the SQDG control (Fig. 6A).

**Figure 6.**
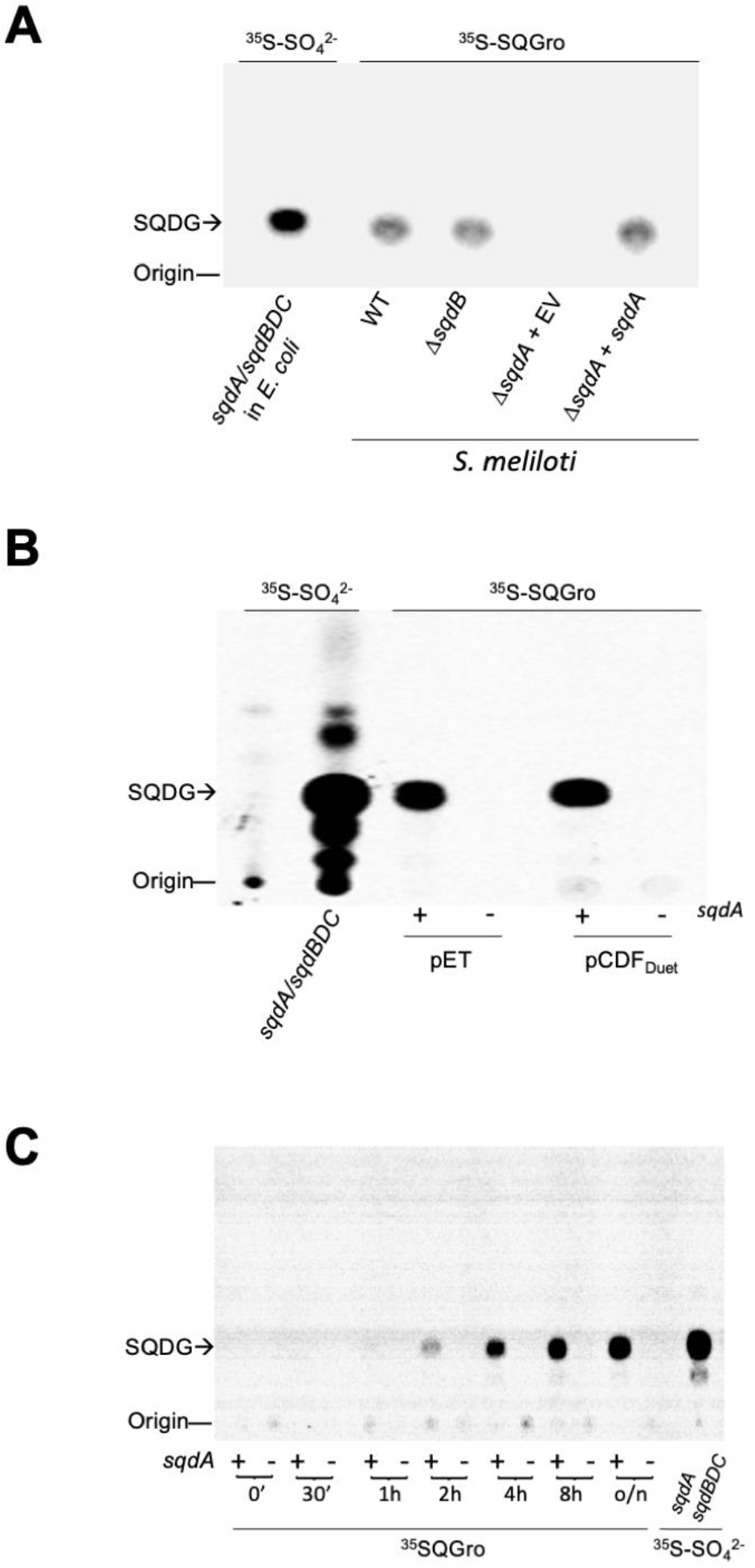
SqdA is required for incorporation of exogenous SQGro into SQDG. **A.** ^35^S-labeled lipids isolated from *S. meliloti* wild type carrying pDRT007 and pDRT008 (*sqdA sqdBDC*), *S. meliloti* wild type (WT), *sqdB* mutant SLD12 (*ΔsqdB*), and *sqdA* mutant NG8 carrying an empty vector (*ΔsqdA* + EV) or complemented with pDRT007 (*ΔsqdA* + *sqdA*) and grown in PYCa^++^ complex medium in the presence of ^35^S-sulfate (*sqdA sqdBDC*) or in the presence of ^35^S-SQGro. **B.** ^35^S-labeled lipids isolated from *E. coli* BL21(DE3) pLysS carrying either pET17b and pRK404 (-), pDRT005 and pDRT008 (*sqdA sqdBDC*), or pET-plasmids (pET) containing *sqdA* (+), or empty vector (-), or pCDF_Duet_ plasmids (pCDF_Duet_) containing *sqdA* (+) or empty vector (-) grown in LB medium in the presence of ^35^S-sulfate or in the presence of ^35^S-SQGro. **C.** Incorporation of ^35^S-SQGro by *E. coli* BL21(DE3) pLysS carrying either the empty plasmid pET17b (-) or the *sqdA* bearing plasmid pDRT005 (+) at the indicated times. As a control SQDG obtained from *E. coli* BL21(DE3) pLysS carrying pDRT005 and pDRT008 (*sqdA sqdBDC*) grown in the presence of ^35^S-labeled sulfate.

Interestingly, SQDG is also formed after addition of SQGro to the *sqdB S. meliloti* mutant which, otherwise, is unable to form SQDG (Weissenmayer *et al*., 2000; Fig. 2). This result indicates that SQGro replaces the function of the *sqdBDC* operon and further supports that SQGro is the product formed by the action of the operon. To convert SQGro to SQDG two acylations are required by at least one acyltransferase (Fig. 1). SqdA is the candidate enzyme to perform at least one of these acylations. Indeed, addition of SQGro to the S. *meliloti* mutant lacking the gene for SqdA did not result in SQDG formation while complementation of such mutant with the *sqdA* gene in *trans* restored formation of SQDG (Fig. 6A). Therefore, SqdA is required to convert exogenous SQGro into the lipid membrane SQDG. Furthermore, addition of SQGro to *E. coli* strains carrying *sqdA* in a pET16b or in a pCDFDuet plasmid leads to SQDG formation while strains carrying the empty plasmid do not form any lipidic ^35^S-labeled compound (Fig. 6B). The formation of SQDG is time dependent and can be observed after 2 hours of induction with IPTG in a *E. coli* strain carrying *sqdA* (Fig. 6C). These results indicate that either SqdA can perform both acylations and converts SQGro into SQDG or enzymes from *E. coli* do the missing acylation reaction. This route of SQDG biosynthesis from SQGro follows the same scheme as the so called glycerophosphocholine (GPC) acylation pathway first described in bacteria in the plant pathogen *Xanthomonas campestris* (Moser *et al*., 2014) and, later, in Gram-positive human pathogens of the streptococci mitis group (Joyce *et al*., 2019). In the GPC acylation pathway, exogenous GPC or lyso-phosphatidylcholine are acylated to form phosphatidylcholine (PC). It was demonstrated that pathogenic streptococci can use the abundant human metabolites GPC and lyso-PC to synthesize the membrane lipid PC (Joyce *et al*., 2019).

SQDGs are very abundant lipids and promiscuous acyl hydrolases convert them sequentially to the metabolites lyso-SQDG and SQGro (Goddard-Borger and Williams, 2017). Recently, different pathways for degradation of sulfoquinovose and SQGro have been described (Sharma *et al*., 2022) that allow to use these compounds as carbon and sulfur source. According to our results, environmental SQGro can also be used as a building block for the synthesis of the membrane lipid SQDG, in an analogous way as human GPC is used as building block for synthesis of PC (Joyce *et al*., 2019).

### SqdA is required for the enzymatic acyl-acyl carrier protein-dependent acylation of lyso-SQDG

Addition of labeled SQGro to an *E. coli* culture carrying *sqdA* led to SQDG formation (Fig. 6), but these *in vivo* experiments did not clarify the specific reaction carried out by SqdA. Previously, we have shown that *S. meliloti* acyltransferases OlsA and OlsB required for ornithine lipid synthesis preferentially use acyl carrier protein (AcpP) as the donor of acyl groups in the respective reactions (Weissenmayer *et al*., 2002; Gao *et al*., 2004). When cell-free extracts of an *E. coli* strain carrying an empty vector were incubated with ^35^S-SQGro and palmitoyl-AcpP, no labeled compounds were detected in the lipidic or in the subsequent *n*-butanol extraction of the aqueous phase of the reaction. However, when the reaction was performed with *E. coli* cell-free extracts that had expressed SqdA, the formation of SQDG as well as SQMG were detected in the lipidic and *n*-butanol phase, respectively (Fig. 7A). When ^35^S-SQMG was used as substrate, the crude extract of an *E. coli* carrying an empty vector could form some SQDG. However, when the *E. coli* extract containing SqdA was used, the amount of SQDG formed was significantly increased (5x). These results clearly show a direct role of SqdA in the acylation of SQMG to form SQDG, and we conclude that SqdA performs the second acylation in SQDG biosynthesis.

**Figure 7.**
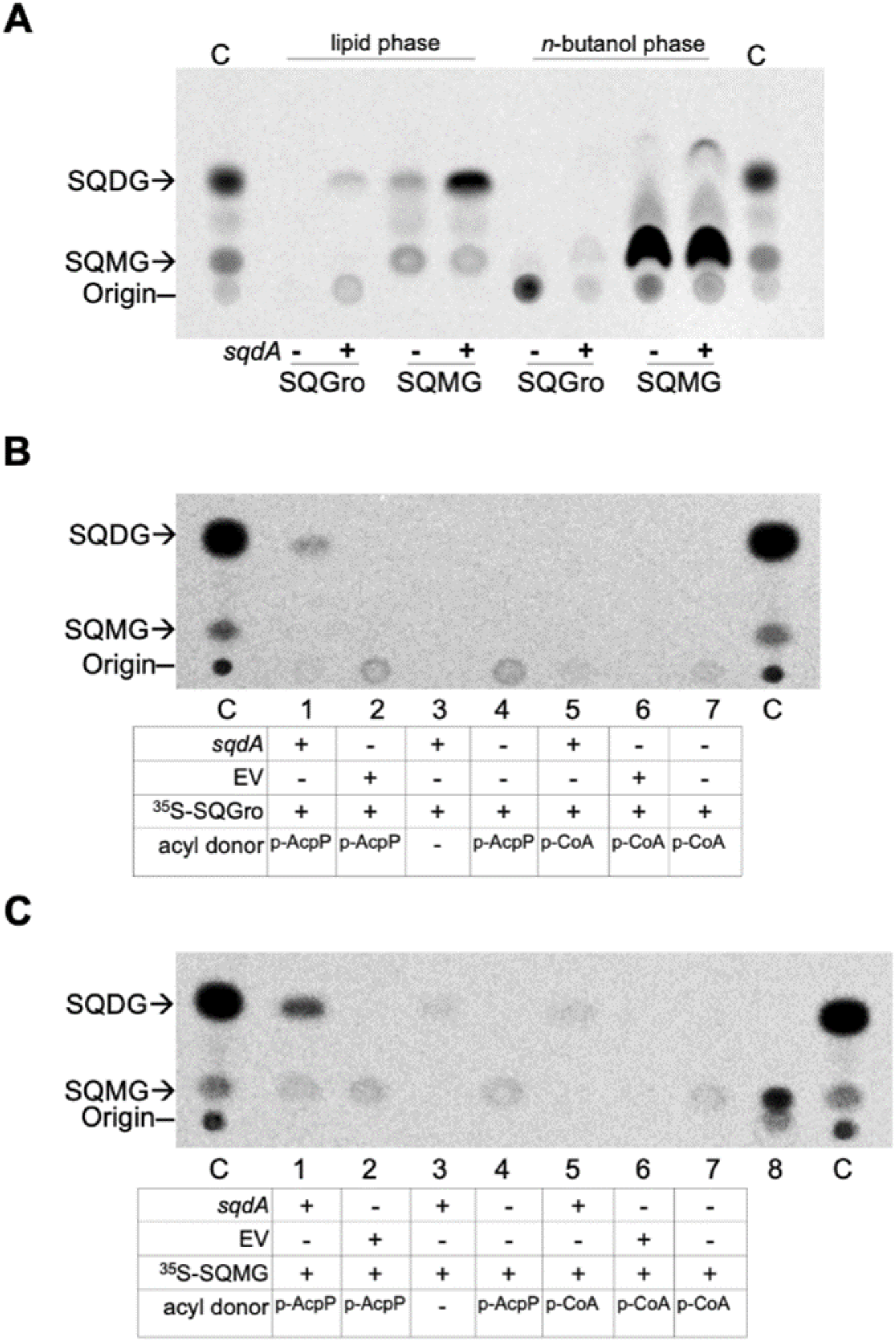
Crude extracts of *E. coli* carrying *sqdA* can convert SQGro and SQMG to SQDG. **A**. Four reactions were set up with crude extracts obtained either from *E. coli* carrying the empty vector pET17b (-) or the *sqdA*-containing plasmid pDRT005 (+). With both extracts the labeled substrates ^35^S-SQGro or ^35^S-SQMG were assayed. All reactions contained palmitoyl-AcpP as acyl donor. Compounds obtained in the chloroform phase of a Bligh and Dyer extraction (lipid phase) are shown. The resulting aqueous phase of the Bligh and Dyer extraction of the reactions was submitted to *n*-butanol extraction (*n*-butanol phase) and separated by one dimension TLC analysis. **B** and **C.** One dimensional TLC analyses of ^35^S-lipids obtained from different *in vitro* reactions using as substrates ^35^S-SQGro (**B**) or ^35^S-SQMG (**C**) and as acyl donors palmitoyl-AcpP (p-AcpP), palmitoyl-CoA (p-CoA) or none (-). The use of crude extracts obtained either from *E. coli* carrying the *sqdA*-containing plasmid pDRT005 (*sqdA*) or carrying the empty vector pET17b (EV) is indicated. In lane 8 of **C**, a portion of the purified ^35^S-SQMG (the same amount as used in reactions) is shown and no SQDG is detectable. Partial hydrolysates of ^35^S-SQDG samples contained SQMG and SQDG and were used as controls (C).

Formation of SQDG from SQGro required the presence of crude extract from *sqdA*-containing strain and palmitoyl-AcpP, while no product was detected when palmitoyl-CoA was used instead of palmitoyl-AcpP (Fig. 7B). For the second acylation step from SQMG to SQDG, some SQDG was formed by SqdA in the absence of acyl donor (Fig 7C, lane 3), that was not increased when palmitoyl-CoA was added to the reaction (Fig. 7C, lane 5). Therefore, SqdA requires acyl-AcpP as donor of the fatty acyl group for the acyltransferase reaction.

### The SQGro acylation pathway appears to be restricted to some members of the

Hyphomicrobiales (Rhizobiales) *and* Rhodobacterales *orders of* Alphaproteobacteria Initial BLASTP searches with the sequences SqdA and SqdD, which are specific for the SQGro acylation pathway (Fig. 1), retrieved only sequences from Alphaproteobacteria. To ascertain how widespread this pathway for SQDG biosynthesis might be, we searched in a collection of 16,076 complete proteobacterial genomes, those carrying ORFs that showed ≥30% protein sequence identity and ≥60% coverage with respect to the *S. meliloti* ORFs. The search returned a total of 238 genomes that encoded the four Sqd proteins. All the genomes were from Alphaproteobacteria and 182 of them belong to the order *Hyphomicrobiales*, that was recently proposed to replace the order *Rhizobiales* (Hördt *et al*., 2020). From those, a total of 160 belong to the family *Rhizobiaceae*, 9 to *Phyllobacteriaceae*, 9 to *Stappiaceae* and 4 to *Aurantimonadaceae.* The other 56 genomes belong to the order *Rhodobacterales* where 48 are from the family *Rhodobacteraceae* and 9 from the family *Roseobacteraceae* (Table S2). Thus, among proteobacteria, the occurrence of the four orthologues for SQDG biosynthesis in one organism occurs in less than 1.5 % of the genomes analyzed. However, since the total number of Alphaproteobacteria present in our collection was 1625, the prevalence of the SQDG acylation pathway in complete genomes amounts to 14.6 % within Alphaproteobacteria.

Villanueva *et al*. (2014) constructed an extended phylogeny of SqdB proteins and one of the clusters they defined, *sqdB2B*, includes only Alphaproteobacteria of the *Hyphomicrobiales* (*Rhizobiales)* and *Rhodobacterales* orders, thus reflecting the distribution of the SQGro acylation pathway. Interestingly, they recovered abundant and diverse gene sequences of the *sqdB2B* cluster in marine microbial mats (Villanueva *et al*., 2014), which might indicate an important prevalence of the SQGro acylation pathway in such an ecosystem.

Presence of only SqdA allows for incorporation of exogenous SQGro into membrane lipids (Fig. 6) and, therefore, we searched for genomes that do not contain SqdB, SqdD or SqdC, but that carried a possibly functional SqdA. We experimentally determined the E value cut-off to be e–40 for a protein to possibly have the function of SqdA and we found 35 genomes that contain SqdA and not any of the other genes for SQDG biosynthesis (Table S3). Most of the genomes that contain an orphan SqdA were of the *Caulobacteraceae* (16) with 7 genomes being *Hyphomicrobiales (Rhizobiales)* and 7 genomes being *Rhodobacterales.* SqdA alone also appears in 3 genomes of the order *Hyphomonadales* and 1 genome of the order *Maricaulares* (Tabla S3). It seems that SqdA has a broader distribution across Alphaproteobacteria than the other genes required for the complete pathway as they are restricted to *Hyphomicrobiales* and *Rhodobacterales* (Table S2).

An evolutionary tree was constructed with representative SqdA sequences (Fig. 8). Sequences of the *Hyphomicrobiales* or the *Rhodobacterales* group among them. Interestingly, in both orders there are genomes that contains orphan SqdAs that show strong relationship to SqdA proteins with demonstrated function (Fig. 8). The prevalence of SqdA alone can provide the organism with the means to scavenge SQGro from the environment to form SQDG as we have proposed. SqdA alone from the order *Caulobacterales*, *Hyphomonadales* and *Maricaulares* appears together in a branch between SqdAs of *Hyphomicrobiales* and of *Rhodobacterales*.

**Figure 8.**
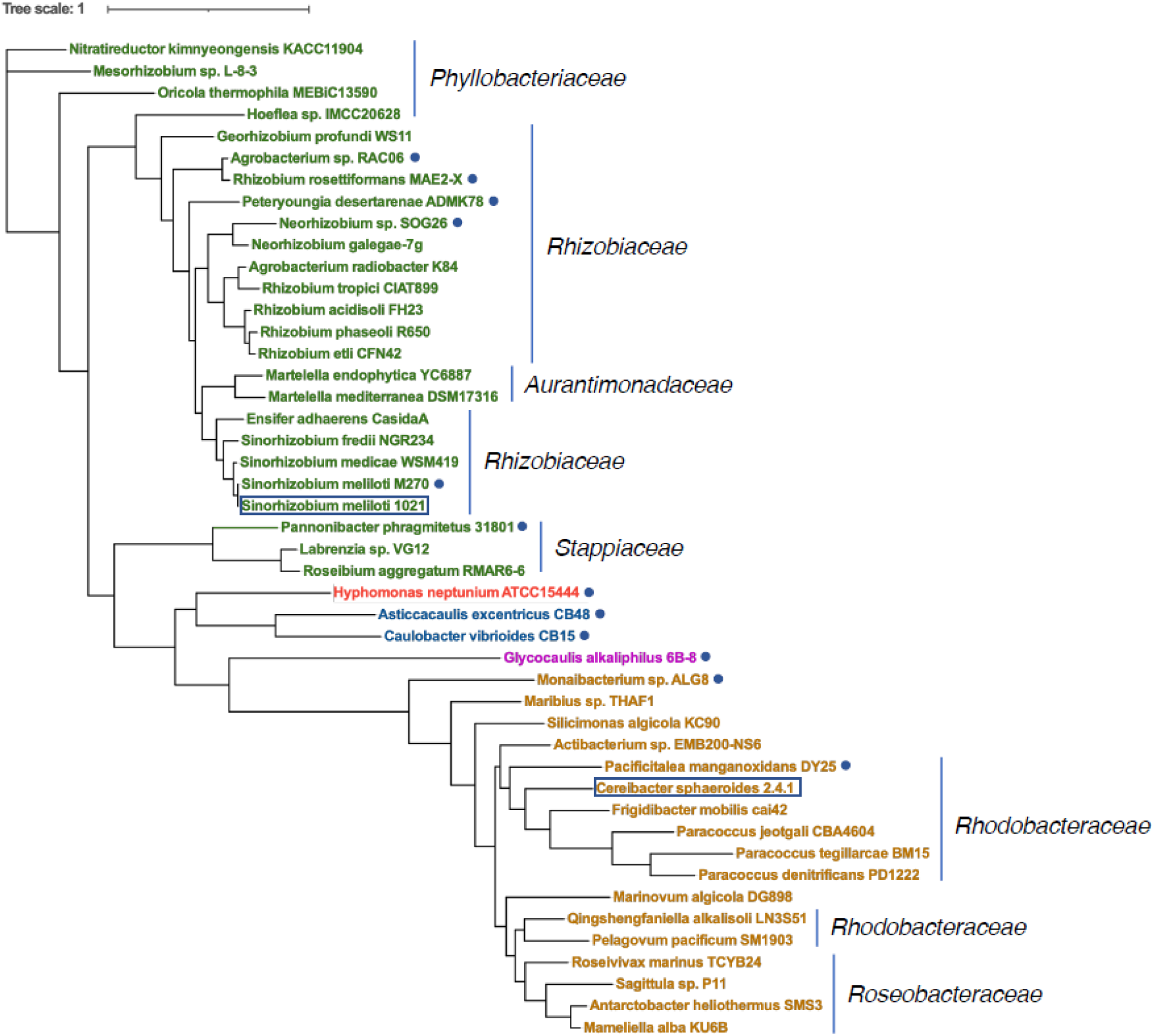
Phylogenetic analysis of the distribution of SqdA proteins. Maximum likelihood phylogenetic tree of homologous proteins from the indicated genomes based on the amino acid sequence of SMc02490. Strains are coloured according to the order they belong to *Hyphomicrobiales* (*Rhizobiales*) in green, *Rhodobacterales* in light brown, *Hyphomonadales* in red, *Caulobacterales* in blue, and *Maricaulares* in pink. Families are indicated next to the tree. Sequences which are encoded in genomes without the other genes for SQDG biosynthesis are marked with a blue dot next to them. The strains *Sinorhizobium meliloti* 1021 *and Cereibacter (Rhodobacter) sphaeroides*, require SqdA for SQDG biosynthesis are enclosed in retancgles. Sequences were aligned using muscle 5.1 (Edgar, 2021). Maximum likelihood phylogenic tree was generated with 100 bootstrap replicates (RAxML 8.2.12). The scale bar is provided as a reference for branch lengths. Gene locus tags are available in Tables S2 and S3.

### SqdA groups with other lyso-lipid acyltransferases that present an HX_5_D motif

Our BLASTP analysis showed a significant similarity between the *S. meliloti* SqdA sequence and the *N-*terminal domain of OlsF proteins (28 % identity and E value of 6e-27 in an overlap of 202 amino acids between SqdA*_Sm_* and OlsF*_Sp_*). OlsF contains two acyltransferase domains that are responsible for the two acylation steps required for ornithine lipid biosynthesis in the Gammaproteobacterium *Serratia proteomaculans* or in *Flavobacterium johnsoniae* (Vences-Gúzman *et al*., 2015). The *C*-terminal domain in OlsF is distantly related to OlsB, an acyltransferase of *S. meliloti* responsible for *N*-acylation of ornithine to form lyso-ornithine lipid (Gao *et al*., 2004). The second acylation of ornithine lipid in *S. meliloti* is carried out by the *O*-acyltransferase OlsA (Weissenmayer *et al*., 2002). Curiously, the *N*-terminal domain of OlsF is unrelated to OlsA, although they both perform acylation of lyso-ornithine lipid (Vences-Guzmán *et al*., 2015; Weissenmayer *et al*., 2002). Heath and Rock (1998) described an HX_4_D motif as indicative for acyltransferases. However, the *N*-terminal domain of OlsF displays an HX_5_D motif (Vences-Guzmán *et al*., 2015). During our alignment to construct the phylogenetic tree of SqdA proteins (Fig. 8) we also found an HX_5_D motif that is strictly conserved in all SqdA sequences. Recently, another lyso-lipid acyltransferase, SalA, with a strictly conserved HX_5_D motif was described. SalA is required for the second acylation step of a newly described sulfur-containing aminolipid (Smith *et al*., 2021). Other well-characterized bacterial lyso-lipid acyltransferases are PlsC, responsible for the final step in the biosynthesis of the phospholipid phosphatidic acid, the aforementioned OlsA, and the lyso-PC acyltransfereses XC_0238 and XC_0188 of *X. campestris* (Moser *et al*., 2014). A phylogenetic tree constructed with different bacterial lyso-lipid acyltransferases clearly separates those having an HX_5_D from those having the HX_4_D motive (Fig. 9). Multiple sequence alignments with representative acyltransferases show the presence of two conserved sequence motifs (Fig. 10), representing the catalytic center (HX_4/5_D) and the possible block III identified in acyltransferases as substrate binding site (Lewin *et al*., 1999), respectively. The presence of the HX_5_D motif in the proteins SqdA, SalA and OlsF deserves further mechanistic analyses.

**Figure 9.**
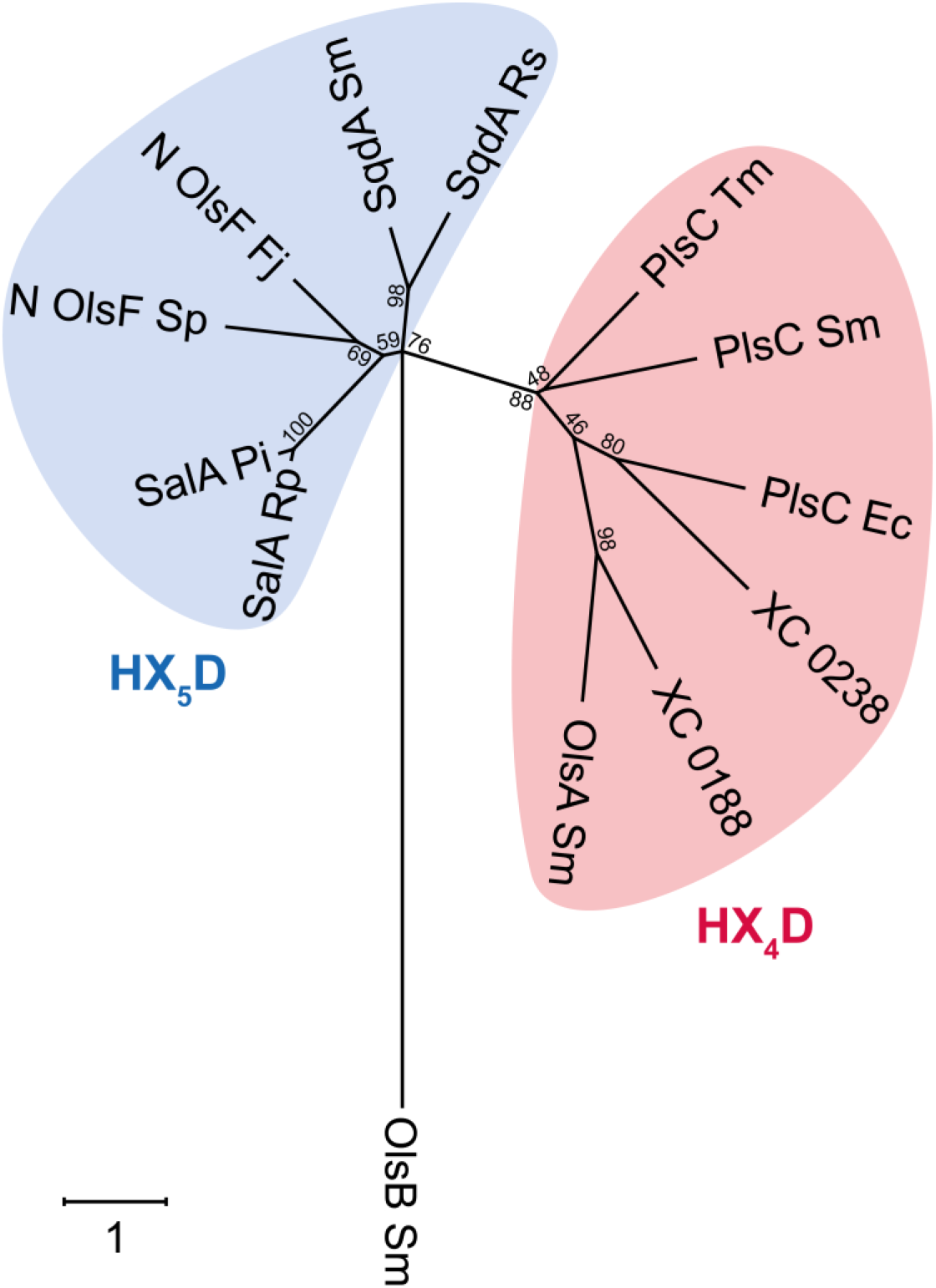
Unrooted phylogenetic tree of selected lyso-lipid acyltransferases with demonstrated function. Amino acid sequences were aligned using the Clustal Omega program (https://www.ebi.ac.uk/Tools/msa/clustalo/). The tree was constructed using the program MEGA version X (http://www.megasoftware.net/) employing the maximum likelihood method. Distances between sequences are expressed as 1 change per amino acid residue. The number at each node represents the bootstrap value as a percentage of 100 replications. ORFs were SqdA from *R. sphaeroides* (SqdA Rs) and *S. meliloti* (SqdA Sm) required for SQDG biosynthesis (Benning and Sommerville, 1992a; this work), *N*-terminal sequences (300 aa) of OlsF sequences from *F. johnsoniae* (N OlsF Fj) and from *S. proteomaculans* (N OlsF Sp) involved in ornithine lipid biosynthesis (Vences-Guzmán *et al*., 2015), SalA from *Phaeobacter inhibens* (SalA Pi) and from *Ruegeria pomeroyi* (SalA Rp) required for biosynthesis of a sulfur-containing amino lipid (Smith et al., 2021); PlsC from *Thermotoga maritima* (PlsC Tm) (Robertson *et al*., 2017), PlsC from *E. coli* (PlsC Ec) and PlsC from *S. meliloti* (PlsC Sm) (Basconcillo *et al*., 2009b), OlsA from *S. meliloti* (OlsA Sm) (Weissenmayer *et al*., 2002), and the two lyso-PC acyltransferases (XC 0238 and XC0188) described in *X. campestris* (Moser *et al*., 2014). The sequence of the *N*-acyltransferase OlsB from *S. meliloti* (OlsB Sm) was used as outgroup (Gao *et al*., 2004).

**Figure 10.**
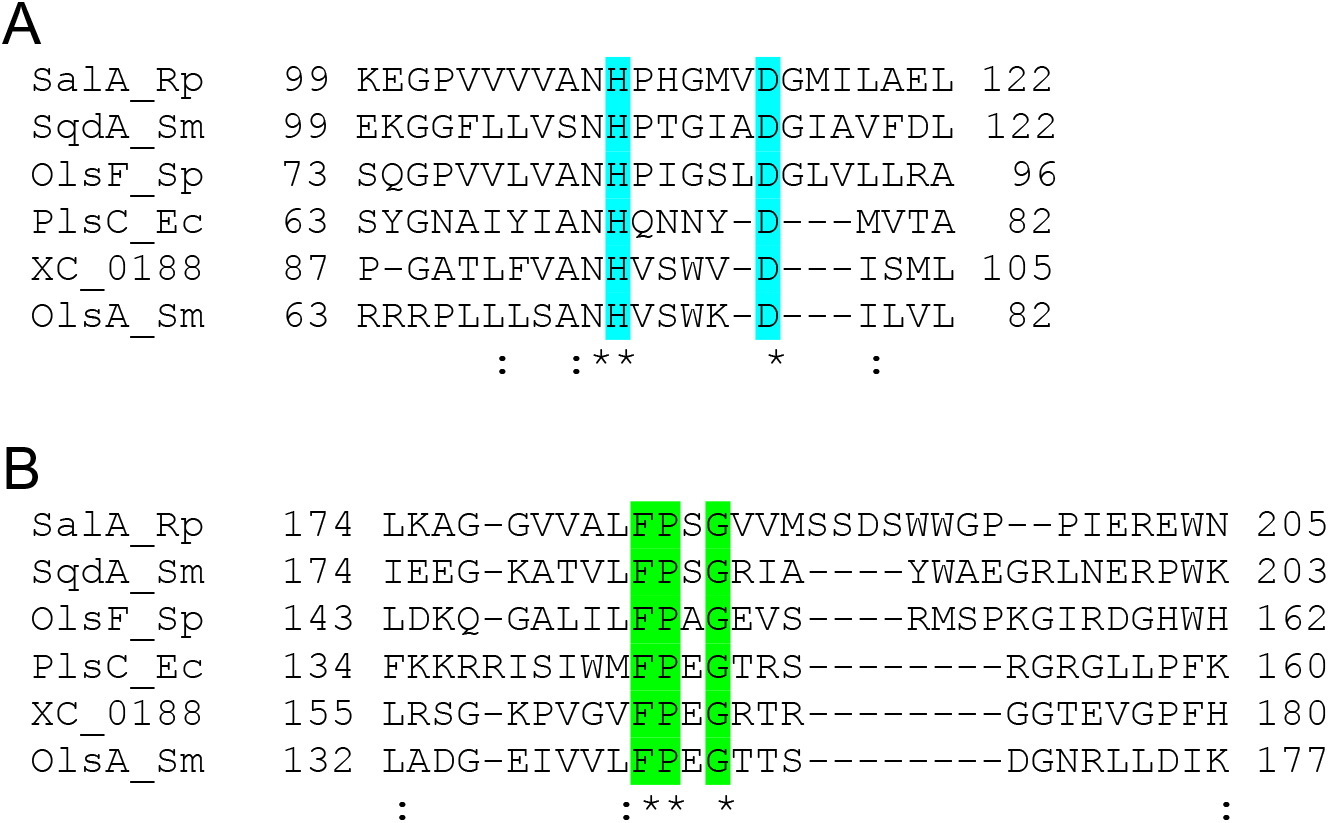
Alignment of two conserved blocks of *S. meliloti* SqdA and other characterized lyso-lipid acyltransferases of bacteria. Amino acid sequences were aligned using the Clustal Omega program (https://www.ebi.ac.uk/Tools/msa/clustalo/). In (**A**) the region containing the amino acids of the catalytic center (HX_4/5_D) with H and D in light blue shading are shown. In SqdA, SalA, and *N*-terminal OlsF there are 5 amino acids between H and D while in PlsC, XC_0188 and OlsA there are 4 amino acids. In (**B**) the region containing the substrate binding site (FPXGXX) with FP and G in green is shown. Description of ORFs are given in figure legend 9.

## Materials and methods

### Bacterial strains, plasmids, and growth conditions

Bacterial strains and plasmids used in this work, as well as their relevant characteristics, are listed in Table 3. Construction of the sinorhizobial mutant deficient in *smc02490 (sqdA)* is described in Table S4. *Sinorhizobium meliloti* strains were cultivated at 30 °C, either in complex medium of yeast extract and peptone (PY) supplemented with 4.5 mM CaCl_2_ (PYCa^++^) (Beringer, 1974) or in Sherwood minimal medium with succinate (8.3 mM) replacing mannitol as the carbon source (Sherwood, 1970), on a gyratory shaker. For determination of lipid composition under phosphate limitation, three subcultivations were carried out in Sherwood minimal medium containing 0.02 mM phosphate. *Escherichia coli* strains were grown at 30 °C in Luria-Bertani (LB) broth (Sambrook and Russell, 2001). Antibiotics for *S. meliloti* were added, when required, at the following final concentrations (μg/ ml) in complex medium: neomycin 200, tetracycline 8, and in liquid media with phosphate limitation, the antibiotic concentrations (μg/ ml) were: neomycin 1 and tetracycline 2. For *E. coli*, antibiotic concentrations were: carbenicillin 100; chloramphenicol 10; tetracycline 10 and kanamycin 50. Plasmids were mobilized into *S. meliloti* by diparental mating using the *E. coli* S17-1 donor strain as previously described (Simon *et al*., 1983).

**Table 3.** Bacterial strains and plasmids used in this study.

| Strain or Plasmid | Relevant characteristics<sup>a</sup> | Reference or source |
| --- | --- | --- |
| *Escherichia coli* |  |  |
| DH5α | *recA1*, Φ80 *lacZ*Δ*MI*; cloning strain | Hanahan, 1983 |
| S17-1 | *thi pro recA hsdR<sup>-</sup> hsdM<sup>+</sup>* RP4 integrated in the chromosome, 2-Tc::Mu, Km::Tn7 (Tp<sup>R</sup>/Sm<sup>R</sup>) | Simon *et al.*, 1983 |
| BL21(DE3) | Expression strain | Studier *et al.*, 1990. |
| *Sinorhizobium meliloti* |  |  |
| *S. meliloti* 1021 | Wild type used throughout this study | López-Lara *et al.*, 2005 |
| our |  |  |
| NG8 | *smc02490::cm* | This study |
| SLD12 | *sqdB::kan* | López-Lara *et al.*, 2005 |
| OD1 | Deficient in OL and DGTS, *olsA::spec*; *btaA::gm* | López-Lara *et al.*, 2005 |
| **Plasmids** |  |  |
| pLysS | Causes repression of T7 polymerase, Cm<sup>R</sup> | Studier, 1991 |
| pET16b | Expression vector, Cb<sup>R</sup> | Novagen |
| pET17b | Expression vector, Cb<sup>R</sup> | Novagen |
| pBBR1MCS-2 | Broad host-range vector, Nm<sup>R</sup> | Kovach *et al.*, 1995 |
| pRK404 | Broad host-range vector, Tc<sup>R</sup> | Scott *et al.*, 2003 |
| pRSFDuet-1 | Expression vector, Km<sup>R</sup> | Novagen |
| pCDFDuet-1 | Expression vector, Sp<sup>R</sup> | Novagen |
| pDRT005 | pET16b carrying *smc02490*, Cb<sup>R</sup> | This study |
| pDRT007 | pBBR1MSC-2 carrying *smc02490*, Nm<sup>R</sup> | This study |
| pBW41 | pUC19a with a 7.2 kb PtsI fragment containing *sqdBDC* | Weissenmayer *et al.*, 2000 |
| pDRT006 | pRK404 with a 7.2 kb PtsI fragment containing *sqdBDC*, Tc<sup>R</sup> | This study |
| pDRT008 | pRK404 carrying *sqdBDC*, Tc<sup>R</sup> | This study |
| pJCR003 | pET17b carrying *sqdB*, Cb<sup>R</sup> | This study |
| pJCR005 | pRSFDuet-1 carrying *sqdB* (mcs-2) and<br>*sqdD* (mcs-1), Km<sup>R</sup> | This study |
| pJCR006 | pRSFDuet-1 carrying *sqdB* (mcs-2) and<br>*sqdDC* (mcs-1), Km<sup>R</sup> | This study |
| pJCR007 | pCDFDuet-1 carrying *smc02490*, Sp<sup>R</sup> | This study |
| pTB5035 | *acpP<sub>Sm</sub>* in pET9a, Km<sup>R</sup> | López-Lara and<br>Geiger, 2000 |
| pAL20 | *acpS<sub>Sm</sub>* in pET9a, Km<sup>R</sup> | Ramos-Vega *et al.*,<br>2009 |
| pAasH | *aas<sub>Ec</sub>* in pET28a, Km<sup>R</sup> | Shanklin, 2000 |

<sup>a</sup>Tc<sup>R</sup>, Tp<sup>R</sup>, Km<sup>R</sup>, Cm<sup>R</sup>, Gm<sup>R</sup>, Cb<sup>R</sup>, Nm<sup>R</sup>: tetracycline, trimethoprim, kanamycine,
chloramphenicol, gentamicin, carbenicillin, and neomycin resistance, respectively.

### DNA manipulations

Recombinant DNA techniques were performed according to standard protocols (Sambrook *et al*., 2001). Commercial sequencing of amplified genes by Eurofins Medigenomix (Martinsried, Germany) corroborated the correct DNA sequences.

Oligonucleotide primer sequences are listed in supplementary Table S5.

### Construction of expression plasmids

The 7.12 kb PstI fragment from pBW41 (Weissemayer *et al*., 2000) carrying the sinorhizobial *sqdBDC* operon was recloned into the broad-host range plasmid pRK404 digested with PstI resulting in plasmid pDRT006, which harbor the *sqdBDC* operon in frame with the *lac* promotor of the vector. After partial digestion of pDRT006 with SphI and subsequent ligation, the plasmid pDRT008 was selected in which the internal 3363 bp SphI fragment of the 7.12 kb sinorhizobial fragment was excised. pDRT008 only contains *sqdBDC* as full ORFs of *S. meliloti*.

Suitable restriction sites for cloning of the respective genes were introduced by PCR in the oligonucleotides (Table S5). Using PCR and specific oligonucleotides oLOP161 and oLOP162, *smc02490* was amplified from *S. meliloti* 1021 genomic DNA. Subsequently, the obtained fragment was cloned into the expression vector pET16b digested with NcoI and BamHI resulting in plasmid pDRT005. The BglII-BamHI fragment of pDRT005 containing *smc02490* was cloned into the BamHI site of pBBR1MSC-2 leading to pDRT007. Using specific oligonucleotides oLOP357 and oLOP358, *sqdB* was amplified by PCR using as template pDRT008. The obtained fragment was cloned into pET17b digested with NdeI and BamHI resulting in plasmid pJCR003. Using the specific oligonucleotides oLOP359 and oLOP360, *sqdD* was amplified by PCR using pDRT008 as template and was cloned in pRSFDuet-1 digested with NcoI and BamHI.

In the resulting plasmid, *sqdB* obtained from pJRC003 as NdeI-BamHI, was cloned into the second multiple site (mcs-2) of pRSFDuet-1 previously digested with NdeI and BglII. The resulting plasmid, containing *sqdB* in mcs-1 and *sqdC* in mcs-2, was named pJCR005. Similarly, *sqdDC* were amplified by PCR from pDRT008 using specific oligonucleotides oLOP359 and oLOP361 and the fragment containing both genes was cloned in pRSFDuet-1 digested with NcoI and BamHI. Afterwards *sqdB* was cloned from pJCR003 into mcs-2 of the pRSFDuet-1 plasmid containing *sqdDC* digested with NdeI and BglII resulting in plasmid pJCR006.

### In vivo labelling of S. meliloti and E. coli with ^14^C-acetate or ^35^S-sulfate and lipid analysis

The lipid compositions of the bacterial strains were determined after labeling with [1-^14^C] acetate (specific activity: 55 mCi/mmol; PerkinElmer) or [^35^S] sulfate (specific activity: 1494 Ci/mmol; PerkinElmer). Cultures of *S. meliloti* (1 ml) were started from precultures grown in the same medium. They were labeled with 1 µCi of [1-14C]-acetate and were incubated for 24 h for the case of growth in PYCa^++^ complex medium or for 48 h in case of growth in minimal medium with low phosphate (0.02 mM). For detection of SQDG and its intermediates after heterologous expression of the *smc02490*, *sqdB*, *sqdC* and *sqdD* genes in *E. coli*, derivatives of the BL21 (DE3) pLysS strain were grown in LB complex medium (1 ml). Cultures were induced at OD_620_= 0.4 with 0.1 mM isopropyl-β-D-thiogalactoside (IPTG) and labeled with 1 µCi of [1-^14^C] acetate or 2 µCi of [^35^S] sulfate, the cultures were incubated for 18 h at 30 °C. To study incorporation of ^35^S-SQGro in *S. meliloti* or *E. coli* cultures, they were grown in 1 ml cultures similarly as described above in the presence of 14000 cpm of ^35^S-SQGro. Cells from bacterial cultures were harvested by centrifugation and resuspended in 100 µl of water. Lipids were extracted according to the method of Bligh and Dyer (1959) and the chloroform or water phases were separated by one-dimensional thin layer chromatography (1D-TLC) on high-performance TLC aluminum sheets (silica gel 60; Merck). For separation of lipids, chloroform: methanol: acetic acid: water (85: 15: 10: 3.5, v / v) was used as the mobile phase. For separation of water-soluble compounds, ethanol: water: acetic acid solvent system (20:10:1, v/v) was used as the mobile phase. ^14^C-Radiolabeled lipids were visualized using a Phosphor-Imager (Storm 820; Molecular Dynamics) and ^35^S-radiolabeled products using a Typhoon FLA 9500. Quantification was performed using ImageQuant TL (Amersham Biosciences).

### Preparation of samples and mass spectrometric analysis

The two *S. meliloti* 1021 strains, the mutant in *smc02490* with an empty vector (NG8 x pBBR1MCS-2) and the mutant supplemented with *smc02490* (NG8 x pDRT007), were grown in 1 L of Sherwood minimal medium with low phosphate (0.02 mM) and corresponding antibiotics and harvested in the third pass after 48 hours of incubation at 30 °C. The strains of *E. coli* BL21 (DE3) pLysS x pET17b x pRK404, *E. coli* BL21 (DE3) pLysS x pDRT005 x pRK404 (expressing *smc02490*), *E. coli* BL21 (DE3) pLysS x pET17b x pDRT008 (expressing *sqdBDC*) and *E. coli* BL21 (DE3) pLysS x pDRT005 x pDRT008 (expressing *smc02490* and *sqdBDC*) were each grown in 500 ml of LB medium with the corresponding antibiotics, induced with 0.1 mM IPTG at an OD_620_ of 0.4 and harvested after 18 h of incubation at 30°C. Lipids from cell pellets were extracted by the Bligh and Dyer method (1959). After redissolving the samples in 200 µL dichloromethane, they were analyzed using MALDI or ESI FT-ICR mass spectrometry in negative mode, using a Bruker Daltonics 9.4 T solariX XR instrument. Analysis of SQDGs from *S. meliloti* and *E. coli* systems: the samples were analysed using MALDI in a matrix of dihydroxybenzoic acid with the following settings: laser power for mass spectral acquisition of 43% (for product ion spectra 48%), 100 shots per scan for mass spectral acquisition (1000 shots per scan for product ion spectra). Each saved spectrum was the sum of 8 scans acquired with 2M data points over an *m/z* range of 300-4000 resulting in a resolving power of ∼200000 at *m/z* 800. External calibration was carried out using sodium trifluoroacetate cluster ions (generated at the same time as the MALDI ions, using the ESI source). Isolation in the quadrupole was with width 2 *m/z* units with fragmentation in the hexapole collision cell achieved using argon gas and a ‘collision energy’ setting of 35 V.

Identification of SQGro: MALDI spectra were generated in a matrix of α-cyano-4-hydroxycinnamic acid under the following conditions: laser power 45%, 200 shots per scan, with each saved spectrum the sum of 16 scans. Spectra were acquired with 2M data points over an *m/z* range of 100-1500, resulting in a resolving power of ∼160000 at *m/z* 300. The spectra were internally calibrated against matrix peaks. ESI mass spectra and product ion spectra were acquired using the following conditions: sample solution was infused at 2 µL/min with a spray voltage of 3500 V and 0.5 s ion accumulation time. Each saved spectrum was the sum of 16 scans acquired with 1M data points over an *m/z* range of 100-1500 resulting in resolving power of ∼80000 at *m/z* 300. External calibration was carried out using sodium trifluoroacetate cluster ions and the spectra were then internally calibrated with peaks from fatty acid background ions.

### SQGro and SQMG preparation

For preparation of ^35^S-labeled SQGro, to a culture of *E. coli* BL21 x pDRT005 x pDRT008 (5 ml) grown in complex LB medium, 0.1mM IPTG and 2 µCi [^35^S]-sulfate were added at an OD_620_ of 0.4 and growth was continued for 18 h. After the culture was collected by centrifugation, total lipids were extracted by the Bligh and Dyer (1959) method. A mild alkali treatment (0.3 M KOH at 37 ° C for 3 h) was applied to the dry lipids to completely cleave the ester bonds. Subsequently, the solution was neutralized with 0.3 M HCl, and hydrophilic components were recovered in the aqueous phase of a Bligh and Dyer (1959) extract. This aqueous extract was separated by one-dimensional TLC using ethanol: water: acetic acid (20:10:1, v/v) as mobile phase. After development, the ^35^S-sulfate labeled SQGro was scraped from the silica plate, extracted 3 times in the aqueous phase of the Bligh and Dyer (1959) solvent system. Non-radiolabeled SQGro was prepared in a similar way from a volume of 30 ml of culture and orcinol dye was used for detection in TLC. Samples were visualized by staining with 0.25 % orcinol in 5% H_2_SO_4_ in ethanol and incubation at 100°C. Glucose was used as standard for quantification of unlabeled SQGro.

The lyso-form of SQDG (SQMG) was preferentially obtained into the water phase of a Bligh and Dyer extract, and we designed a protocol to obtain purified SQMG that was not containing SQGro or SQDG. ^35^S-labeled-lipids were obtained by Bligh and Dyer extraction from *E. coli* BL21(DE3) pLysS pDRT005 pDRT008 grown as explained above. A partial mild alkali hydrolysis was achieved by treating lipids with 3 mM KOH for 45 min at 37 °C and then neutralized with 3 mM HCl. The mixture was subjeced to Bligh and Dyer extraction and the water phase was recovered. After two extractions with *n*-butanol of this water phase, purified SQMG is obtained. ^35^S-SQGro and ^35^S-SQMG were quantified by multi-purpose scintillation counter Beckman Coulter LS 6500.

### Preparation of palmitoyl-AcpP and in vitro acylation assays with SQGro or SQMG

Purified apo-AcpP of *S. meliloti* was obtained from *E. coli* BL21 (DE3) pLysS containing plasmid pTB5035 and was converted to functional holo-AcpP using holo-ACP synthase from *S. meliloti* as previously described (Ramos-Vega *et al*., 2009). To obtain palmitoyl-AcpP, holo-AcpP was incubated with palmitic acid (PO500, Sigma) in the presence of the *E. coli* inner membrane enzyme acyl-ACP synthetase (Aas) obtained from the *E. coli* BL21(DE3) pLysS strain carrying the plasmid pAasH (Shanklin, 2000). The palmitoyl-AcpP obtained was purified by DEAE-Sepharose chromatography. *E. coli* BL21 (DE3) pLysS cells carrying the empty vector pET17b or plasmid pDRT005 (*sqdA*) were grown on 30 ml of LB medium with chloramphenicol and carbenicillin from overnight inocula grown in the same medium. Cultures were induced at OD_620_= 0.4 with 0.1 mM IPTG and further grown for 4 h at 30 °C. Cells were harvested, resuspended in 1 ml of 100 mM Tris-HCl pH 8.5 buffer and kept frozen at - 20 °C. Cells were broken by 3-4 steps of freezing and thawing. Afterwards, 25 units of DNAase I and Mg^2+^ to 5 mM final concentration were added and extract were incubated for 20 min at room temperature. The lysates were centrifuged at 7000 g for 20 min at 4 °C and the supernatants were used as cell-free extracts.

Each *in vitro* reaction contained 20 µl of the respective cell-free extract, 14000 cpm of ^35^S-SQGro or ^35^S-SQMG (or 10000 cpm in the case of experiments in Fig. 7B and C), 100 mM Tris-HCl pH 8.5 and 40 µM palmitoyl-AcpP or palmitoyl-CoA (P9716 Sigma). Reactions were incubated at 30 °C for 4 h and lipids were extracted by Bligh and Dyer. Then, the aqueous phases were submitted to *n*-butanol extraction to recover SQMG. Lipids were analyzed by one-dimensional TLC and phosphorimaging as described above. A ^35^S-SQDG sample submitted to partial mild alkaline hydrolysis served as standard for SQDG and SQMG.

### Bioinformatic analyses

Phylogenetic analysis to search for the presence in genomes of genes for SQDG biosynthesis was done by separate BLASTP searches of the *S. meliloti* 1021 SMc02490 (SqdA), SqdB, SqdD and SdqC proteins against 16,076 complete proteobacterial genomes from GenBank that had identical GCA (GenBank assemblies) and GCF (RefSeq assemblies) annotations. Homologues to each ORF with at least 60 % sequence coverage and 30 % amino acid identity to the query sequences were included in the search (Table S2). SqdA sequences were aligned using muscle 5.1 (Edgar, 2021). The evolution model was calculated with ProtTest3 (Darriba *et al*., 2011). The phylogenetic analysis was done RAxML 8.2.12 (Stamatakis, 2014).

## Author contributions

JYCR, NGS, EB, JTO, OG, and IMLL designed the study. JYCR, DRT, NGS, AMO, GG, EB, and IMLL carried out the experiments. All authors carried out the data analysis and discussed the results. JYCR, JTO, OG and IMLL, were involved in drafting the manuscript and all authors read and approved the final manuscript.

## Acknowledgements and funding

J.Y. C-R. is a doctoral student from Programa de Doctorado en Ciencias Biomédicas, Universidad Nacional Autónoma de México (UNAM) and received fellowship 596234 from CONACyT. We acknowledge Jonathan Padilla-Gómez for preparing Figure 9. We thank Lourdes Martínez-Aguilar and Miguel Ángel Vences-Guzmán for skillful technical assistance. This work was supported by projects UNAM-PAPIIT (IN202616 and IN200819) as well as by CONACyT 153998 to IM L.-L. OG acknowledges funding from UNAM-PAPIIT (IN201120). The mass spectrometric analyses were carried out using one of the instruments in the University of York Centre of Excellence in Mass Spectrometry, which was created thanks to a major capital investment through Science City York, supported by Yorkshire Forward with funds from the Northern Way Initiative, and subsequent support from EPSRC (EP/K039660/1; EP/M028127/1).

## Supplementary Information

**Table S1.** Membrane lipid composition of *Sinorhizobium meliloti* 1021 carrying either two empty vectors (1021 x pBBR1MCS-2 x pRK404), *smc02490* in pBBR1MCS-2 and pRK404 vector (1021 x pDRT007 x pRK404), pBBR1MCS-2 vector and *sqdBDC* in pRK404 (1021 x pBBR1MCS-2 x pDRT008) or *smc02490* in pBBR1MCS-2 and *sqdBDC* in pRK404 (1021 x pDRT007 x pDRT008) after growth on low (0.02 mM) phosphate-containing minimal medium or on PYCa^++^ complex medium.^a^

| Composition (% of total $^{14}\text{C}$) | | | | | | | | |
| --- | --- | --- | --- | --- | --- | --- | --- | --- |
|  | 1021 x pBBR1MCS-2 x pRK404 |  | 1021 x pDRT007 x pRK404 |  | 1021 x pBBR1MCS-2 x pDRT008 |  | 1021 x pDRT007 x pDRT008 |  |
| Lipid | low phosphate | complex medium | low phosphate | complex medium | low phosphate | complex médium | low phosphate | complex medium |
| DGTS | 62.3 ± 1.6 | ND | 53.8 ± 2.4 | ND | 59.1 ± 1.2 | ND | 43.3 ± 0.2 | ND |
| MMPE + CL | 6.8 ± 1.2 | 30.7 ± 2.3 | 7.3 ± 1.5 | 28.3 ± 0.9 | 7.0 ± 1.2 | 28 ± 3.8 | 9.3 ± 0.8 | 27.5 ± 4.7 |
| PE+ OL | 12.3 ± 1.6 | 19.7 ± 0.1 | 8.8 ± 3.2 | 19.7 ± 1.1 | 11.1 ± 1 | 19.5 ± 2.2 | 9.7 ± 2 | 19.2 ± 4.4 |
| PG | 5.8 ± 1 | 10.4 ± 1.6 | 3.5 ± 0.6 | 9.1 ± 1.1 | 6.9 ± 0.8 | 9.5 ± 0.8 | 3.6 ± 0.3 | 7 ± 0.8 |
| PC | 4.5 ± 0.9 | 37 ± 1.1 | 6.5 ± 1.4 | 40 ± 1.3 | 5.7 ± 0.7 | 39.7 ± 5.2 | 10.6 ± 1.4 | 37.8 ± 11.2 |
| SQDG | 8.3 ± 1.6 | 2.2 ± 0.2 | 20.1 ± 2.6 | 2.9 ± 0.9 | 10.2 ± 2.8 | 3.3 ± 0.4 | 23.5 ± 3.4 | 8.5 ± 2.9 |

<sup>a</sup> The values shown are mean values ± standard deviation derived from three independent experiments. ND, not detected.

**Table S2.** Alphaproteobacteria containing the four ORFs SqdA, SqdB, SqdD and SqdC for SQDG biosynthesis. For each ORF, an identity ≥ 30 % and a coverage of at least 60 % to the *S. meliloti* 1021 ORFs was considered. (Excel document Table S2).

**Table S3.** Alphaproteobacteria containing a copy of SqdA showing an E value of less than e-40 and which do not contain homologous for SdqB, SqdD or SqdC. (Excel document Table S3).

**Table S4.**
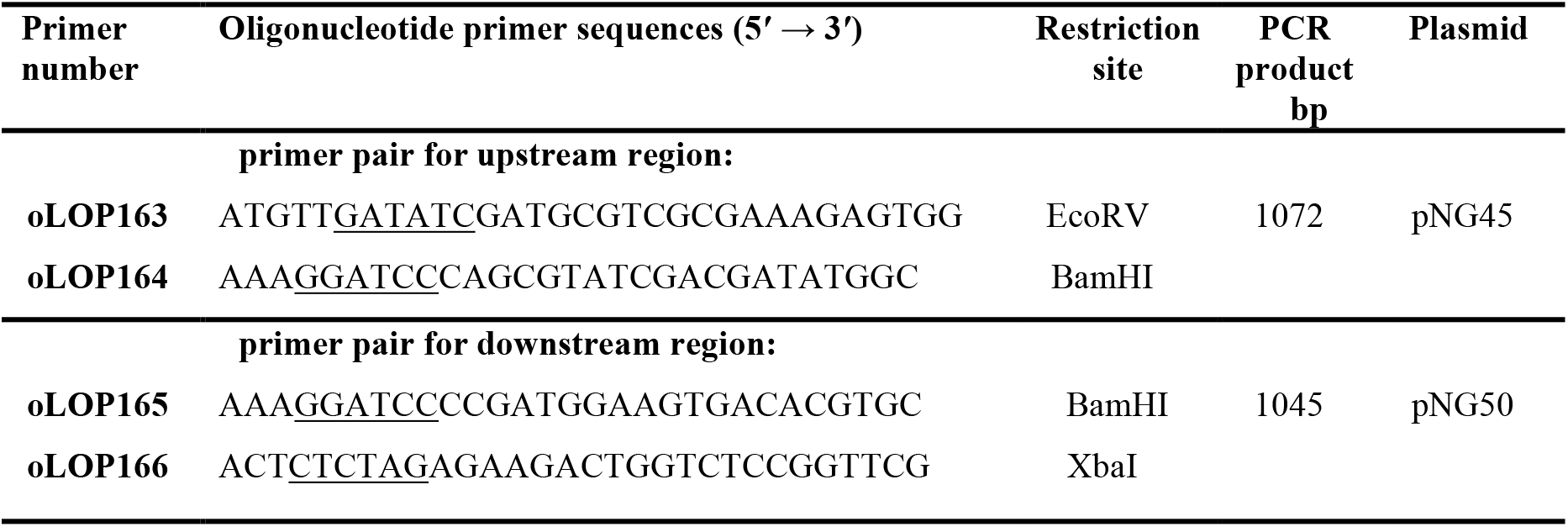
Oligonucleotides primers used for construction of NG8 mutant (Δ*smc02490*). Introduced restriction sites are underlined. For inactivation of the gene coding for SMc02490, the upstream and downstream regions were PCR-amplified (XL-PCR kit; Applied Biosystems) with each specific pair of oligonucleotide primers from genomic DNA of *S. meliloti* 1021, introducing restriction sites (underlined). Subsequently, the upstream region was cloned into the pCR2.1-TOPO® vector (Invitrogen) and the downstream region into pBluescriptSK + (Stratagene) resulting in plasmid pNG45 and pNG50, respectively. The 1072 bp EcoRV-BamHI upstream fragment was excised from pNG45 and was cloned into pNG50 digested with EcoRV-BamHI, resulting in plasmid pNG52. Then, the chloramphenicol resistance-conferring cassette was obtained from pCAT (González-Silva *et al*., 2011) as BamHI fragment and cloned between upstream and downstream regions contained in pNG52. From here, the regions usually flanking *smc02490* and the chloramphenicol resistance-conferring cassette located in between those regions were cloned into the suicide vector pK18*mobsacB* (Schäfer *et al*., 1994) leading to plasmid pNG54. The construct was introduced into wild type *S. meliloti* 1021, and double recombinants in which chloramphenicol resistance-conferring cassette had replaced the sinorhizobial gene were obtained following a procedure described previously (Sohlenkamp *et al*., 2004). Mutations in which the original sinorhizobial gene was replaced by the chloramphenicol resistance-conferring cassette were confirmed by Southern-blot hybridization.

**Table S5.**
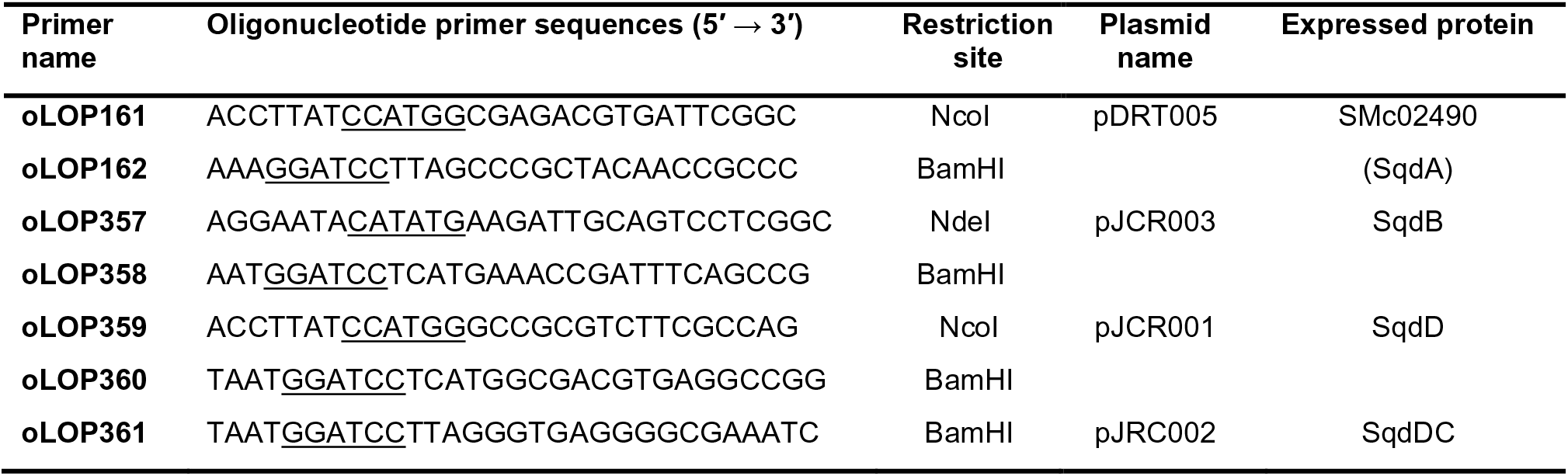
Oligonucleotides primers used for construction of expression plasmids.

---

**Supplementary figure S1.**
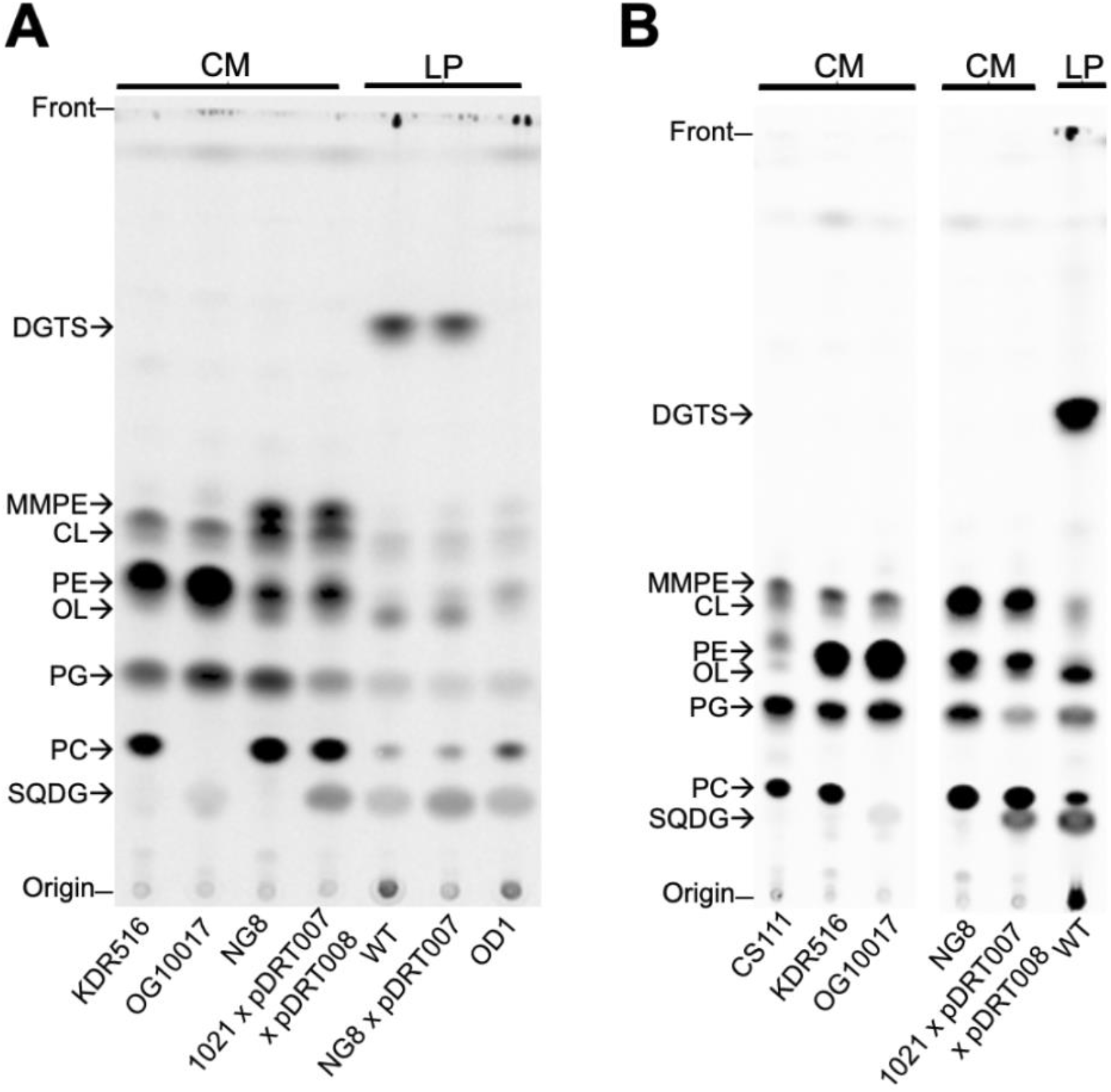
Lipid profile of different *S. meliloti* strains resolved by one-dimensional TLC. **A** and **B** are two different examples of TLCs using ^14^C-acetate labeled lipids obtained from *S. meliloti* strains grown either for 24 hours on complex media (CM) or for 48 hours on low (0.02 mM) phosphate-containing minimal medium (LP). Lipids were obtained from the strains KDR516 which lacks MMPE and DMPE (de Rudder *et al*., 2000), OG10017 which lacks MMPE, DMPE and PC (de Rudder *et al*., 2000), NG8 which lacks SQDG (this work). The wild type 1021 containing plasmids pDRT007 and pDRT008, increased production of SQDG on CM (this work) whereas wild type 1021 (WT) grown on LP forms DGTS, OL and SQDG major lipids (Geiger *et al*., 1999). NG8 carrying pDRT007, increased the amount of SQDG (this work), OD1, lacks DGTS and OL (López-Lara *et al*., 2005), and CS111, lacks PE (Sohlenkamp *et al*., 2004). Spots for PE and OL were not separated and, therefore, were quantified together. In some cases, as in (**B**), CL and MMPE were not completely separated and were quantified together. In the examples shown here, the complex medium used was LB/MC containing 10 mM CaCl_2_ that confers similar growth rate to strains 1021, CS111 and OG10017 (Geiger *et al*., 2021).

**Supplementary figure S2.**
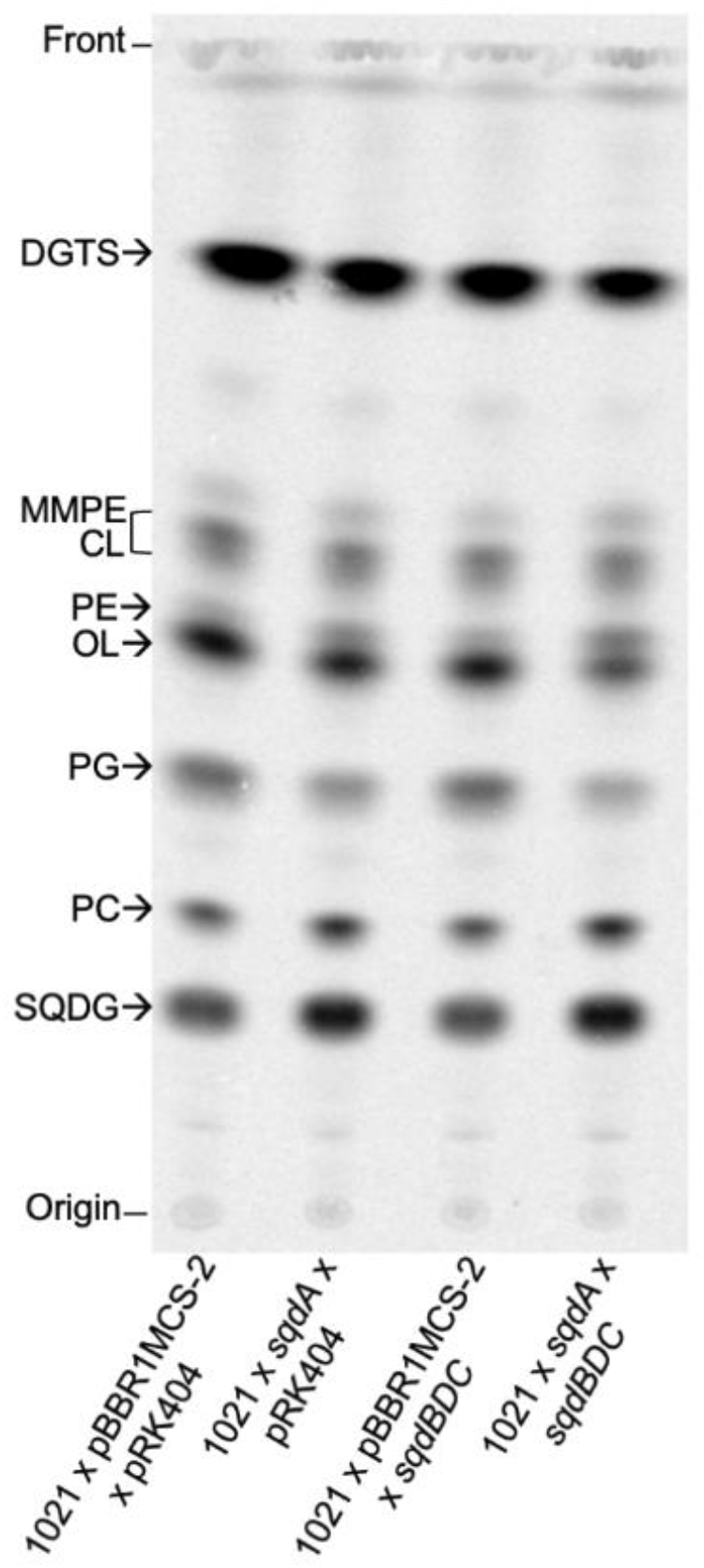
Separation on TLC of ^14^C-labeled lipids formed by *S. melioti* 1021 wild type carrying either two empty vectors (1021 x pBBR1MCS-2 x pRK404), pDRT007 and pRK404 (1021 x *sqdA* x pRK404), pDRT008 or pBBR1MCS-2 (1021 x *sqdBDC* x pBBR1MCS-2) or pDRT007 and pDRT008 (1021 x *sqdA* x *sqdBDC*) grown on Sherwood minimal medium with limiting (0.02 mM) concentrations of phosphate. A representative TLC out of three is shown and the corresponding lipid quantifications are presented in Table S1.

**Supplementary figure S3.**
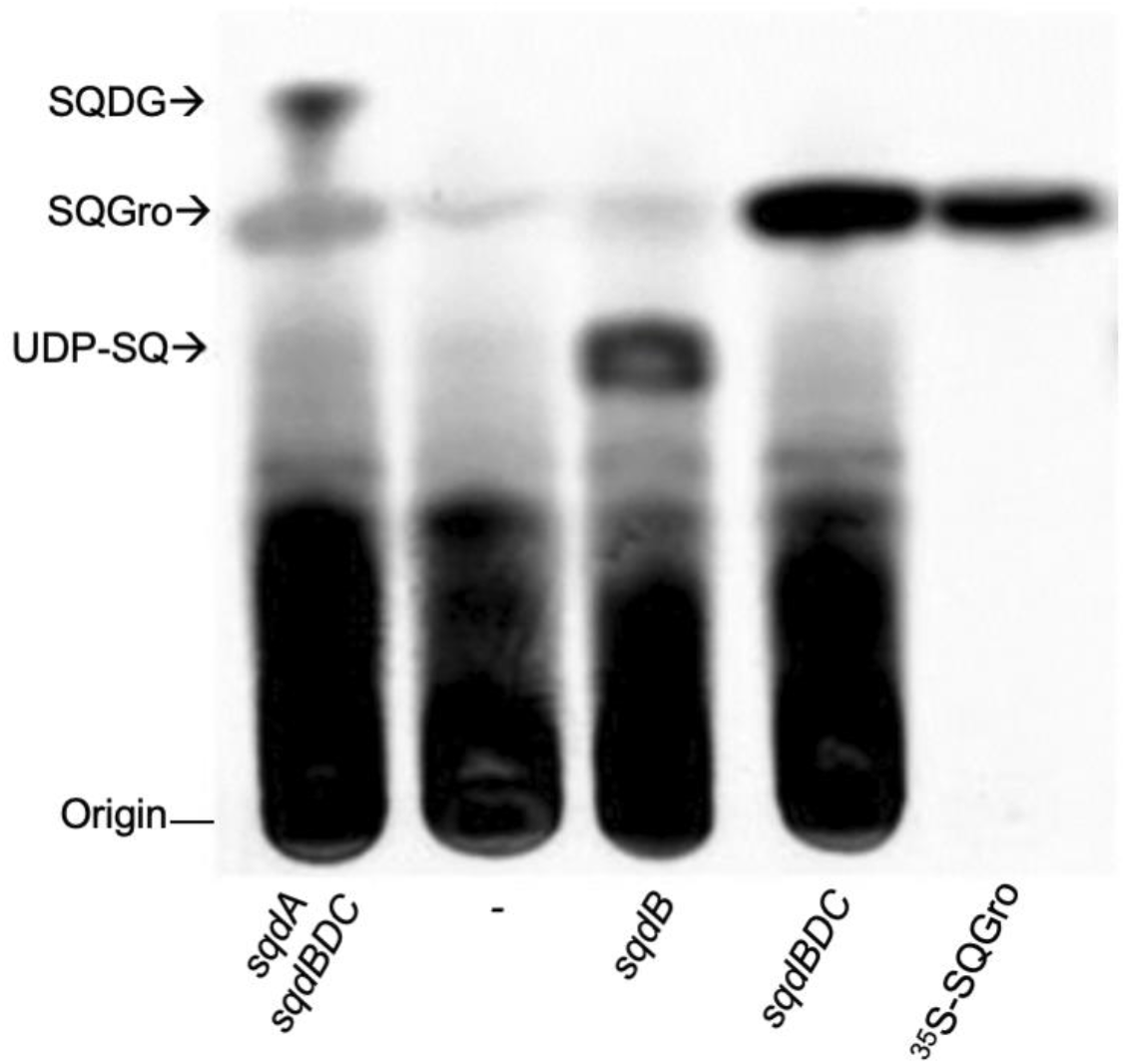
SQGro is an intermediary in the sinorhizobial pathway for SQDG biosynthesis. Thin layer chromatography of ^35^S-sulfate labelled, water-soluble compounds obtained from *E. coli* BL21(DE3) pLysS carrying either pDRT005 and pDRT008 (*sqdA sqdBDC*), the empty vector pET17b (-), pJCR003 (*sqdB*), pJCR006 (*sqdBDC*_Duet_), in comparison to ^35^S-labeled sulfoquinovosyl glycerol prepared from purified ^35^S-SQDG (^35^S-SQro). UDP-SQ: UDP sulfoquinovose. The mobile phase was ethanol: water: acetic acid (20:10:1, v/v).

